# Speciesformer learns conserved cellular states for cross-species generative virtual cell modeling

**DOI:** 10.64898/2026.09.22.752128

**Authors:** Jiacheng Wang, Jiaqi Dong, Guowei Li, Chao Fang, Liwei Liu, Xin Gao

## Abstract

Cells across species are governed by evolutionarily conserved biological programs, yet their molecular states and responses are reshaped by species, tissue and cellular context. A central challenge for virtual cell modeling is therefore to learn cellular states and state transitions that separate transferable biological principles from context-specific variation. Existing single-cell foundation models have advanced cellular representation learning, but most remain focused on single-species analysis, discriminative tasks or specialized forms of generation. Here we present Speciesformer, a cross-species generative single-cell foundation model that integrates evolutionary representation learning with virtual cell-state generation. Speciesformer is pretrained on SpeciesCorpus, comprising 131 million cells from 11 species, 154 tissues, and more than 923 cell types, and maps species-specific genes into a shared evolution-informed gene space. Its encoder learns transferable cell and gene representations that support biological representation probing and cross-species knowledge transfer, providing a common state space for resolving conserved and context-dependent cellular programs. Building on this shared representation, Speciesformer uses a unified generative architecture to model cellular states and state transitions under semantic and interventional conditions, enabling bidirectional generation between transcriptomic states and biological text descriptions, as well as prediction of post-perturbation transcriptomes from initial cell states and intervention descriptions, including in previously unobserved cellular contexts. By unifying evolutionary variation, biological semantics and conditional state transitions, Speciesformer extends cross-species foundation modeling toward a generative virtual cell framework for representing, describing and predicting cellular states across biological contexts.

## 1. Introduction

Single-cell sequencing has made it possible to profile cellular identity and state at unprecedented scale across tissues, developmental stages, diseases and species (Ding, et al., 2022). These data reveal that cellular phenotypes arise from a combination of broadly conserved regulatory programs and context-dependent variation shaped by lineage, tissue environment and physiological state. A central goal of virtual cell modeling is therefore not simply to reproduce observed transcriptomes, but to learn a representation of cellular states and state transitions that separates general biological rules from context-specific variation (Bunne, et al., 2024). Such a model should capture what is shared across biological systems, recognize when those rules diverge, and use this structure to describe, generate and predict cellular states under new conditions. Cross-species comparative biology provides a natural framework for learning this distinction. Evolution acts over long timescales to preserve some cellular programs while rewiring others, creating a diverse set of biological contexts in which conserved and lineage-specific mechanisms can be observed. Comparing cells across species can therefore help distinguish stable cellular programs from species-specific expression patterns and provide a stronger test of biological generalization than within-species interpolation alone. The rapidly expanding collection of single-cell atlases across the tree of life now makes it possible to learn shared representations across evolution, transfer knowledge from data-rich to data-poor organisms and study how cellular functions are conserved or modified across biological contexts (Abdulla, et al., 2024)(Rood, et al., 2025)(Edgar, et al., 2002). This evolutionary diversity offers both a source of biological variation and an inductive bias for identifying transferable cellular principles.

Single-cell foundation models have emerged as a promising approach for learning reusable cell and gene representations from large transcriptomic corpora (Theodoris, et al., 2023)(Cui, et al., 2024)(Hao, et al., 2024)(Pearce, et al., 2026). Recent cross-species models further show that evolution-aware representations can support cell alignment, comparative analysis and knowledge transfer across organisms (Tarashansky, et al., 2021)(Yang, et al., 2024)(Rosen, et al., 2026)(Pearce, et al., 2026). However, most current approaches remain centered on encoding observed cells for tasks such as annotation, retrieval, integration or regulatory analysis. These capabilities are important, but they address only one part of virtual cell modeling. A virtual cell model should also connect molecular states to interpretable biological semantics and generate cellular states that are not directly observed in the training data. This requires moving from representation learning toward generative modeling of both cellular identity and state transitions. Biological descriptions offer a flexible representation of cell type, functional state, tissue context and experimental condition. Mapping expression profiles to such descriptions can make learned cell representations more interpretable, while the reverse modeling allows biological descriptions to specify cellular states that can be generated in expression space. Perturbation modeling extends this problem from state generation to state transition generation. CRISPR-mediated gene knockout (Xue, et al., 2014), RNA interference (Kamath, et al., 2003), transcription factor modulation (Gilbert, et al., 2014), small-molecule treatment (Srivatsan, et al., 2020), cytokine stimulation (Schmidt, et al., 2022), morphogen exposure(Azbukina, et al., 2026) and other environmental or developmental interventions alter cellular programs in a context-dependent manner, and the same perturbation can produce different outcomes across cell types, tissues, disease states or other biological settings. A useful virtual cell model should therefore not only reconstruct observed perturbation responses, but also transfer perturbation dynamics to cellular contexts that have not been experimentally profiled.

In this study, we present Speciesformer, a cross-species single-cell generative model designed as a step toward this broader virtual cell objective. Speciesformer is pretrained on SpeciesCorpus, comprising approximately 131 million cells from 11 species, retrieved and harmonized from CELLxGENE (Abdulla, et al., 2024) and GEO (Edgar, et al., 2002). To enable cross-species modeling without relying on one-to-one orthologous gene mapping, we developed an evolution-informed macrogene vocabulary to map species-specific genes into a shared gene space. This strategy allows the model to compare and integrate expression programs across evolutionarily distant organisms while preserving species-specific transcriptional information. Speciesformer learns transferable cell and gene representations for biological probing, including cell type annotation, genetic perturbation modeling and gene-level regulatory analysis, while also supporting cross-species cell alignment and knowledge transfer. Building on this shared cellular state space, Speciesformer further connects transcriptomic representations with biological descriptions and enables bidirectional generation between cell expression and biological semantics. The same generative framework is adapted to model perturbation-induced transitions from an initial cell state to an intervention-conditioned state. By integrating evolutionary variation, biological semantics and conditional state generation within a unified architecture, Speciesformer aims to model both conserved cellular principles and context-dependent responses, moving cross-species foundation modeling toward a generative virtual cell framework.

Our results show that the SpeciesCorpus and the unified generative framework enables the model to transfer context-specific and perturbation-specific cellular states, including combinations of cell types, biological contexts or perturbation conditions not directly observed during training. Notably, the model exhibits strong zero-shot generalization, achieving performance comparable to fine-tuned baselines on benchmark tasks. After task-specific fine-tuning, it further substantially outperforms existing baseline models across diverse generative benchmarks. These capabilities demonstrate that a cross-species generative framework can unify cellular representation learning, comparative cell-state alignment, semantic generation and perturbation response prediction, providing a general framework for virtual cell modeling across species.

## Results

### Speciesformer a unify foundation model for multi-species encoding and generating

We sought a unified model that learns conserved and divergent rules of gene expression across species, and that can generate virtual cells conditioned on text prompts describing cell identity or perturbations (Fig. 1a). To this end, we assembled SpeciesCorpus, a large-scale multi-species single-cell pretraining corpus comprising 131 million cells from 11 species, 462 datasets and 154 tissues (Fig. 1b, Supplementary Table 1). We represented genes using ESM-2 (Lin, et al., 2023) protein embeddings, but instead of treating these embeddings directly as gene tokens, as in existing cross-species models (Rosen, et al., 2026)(Pearce, et al., 2026), we used them to construct a *Deuterostomia*-level macrogene vocabulary through an inverse evolutionary-tree strategy (Fig. 1c). This design maps species-specific genes into a shared gene space while preserving evolutionary structure. Each cell is accompanied by rich cell-level metadata and gene-level priors, enabling the model to learn cellular programs across multiple biological scales.

**Figure 1.**
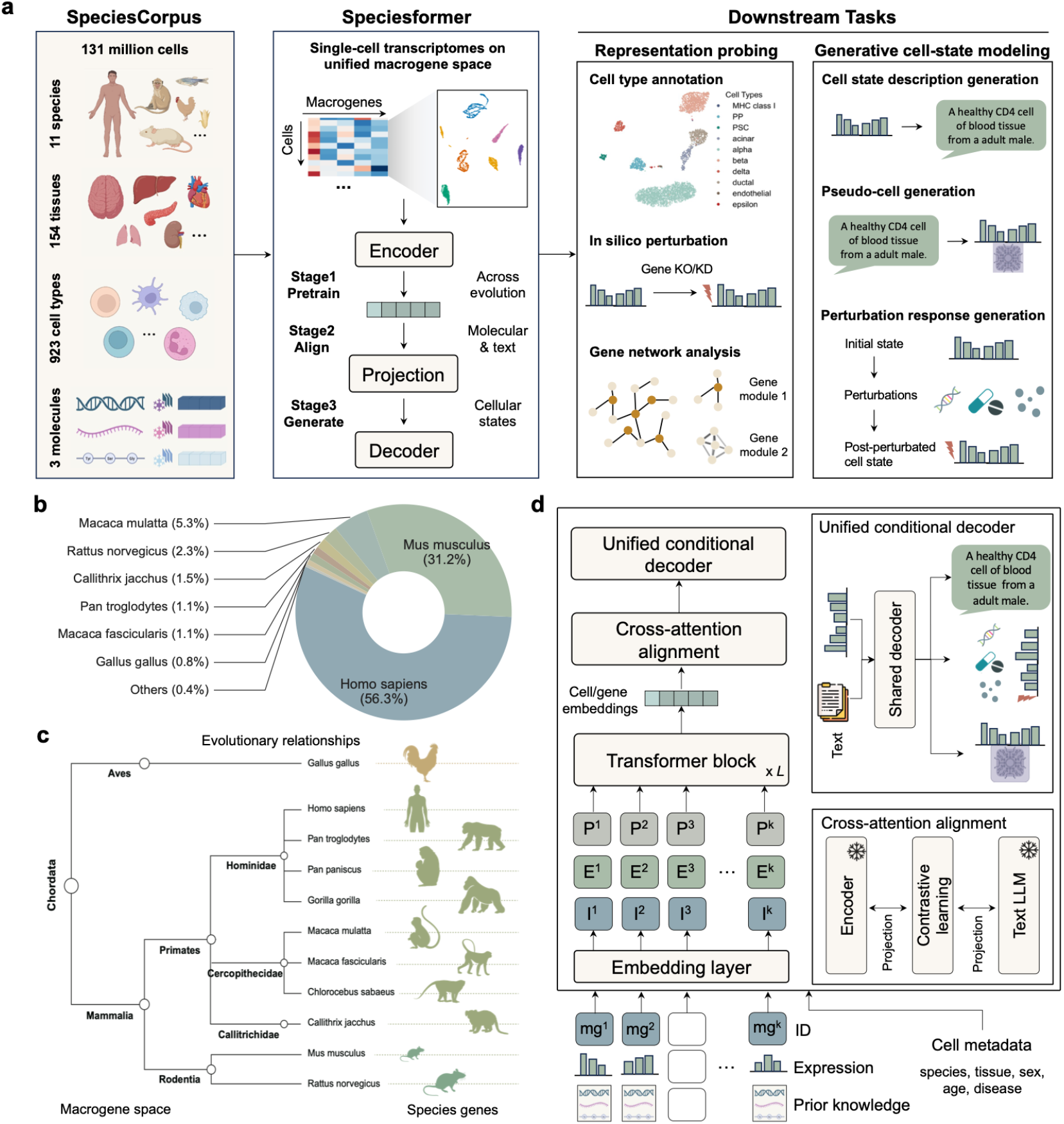
Overview of Speciesformer. **a**, Schematic of Speciesformer. Speciesformer was pretrained on SpeciesCorpus, containing approximately 131 million cells across 154 tissues and 923 cell types. Multi-species single-cell transcriptomes are represented in a unified macrogene space and encoded into cell-level and gene-level representations. These cellular and molecular modalities are aligned with biological text by a projection. The cross-species encoder learns transferable cell and gene embeddings for downstream biological analyses, while a unified generative decoder support cell-state description generation, pseudo-cell expression generation and perturbation-conditioned expression generation. **b**, Species composition of SpeciesCorpus. **c**, Evolutionary structure of the pretraining species, supporting the construction of a shared macrogene vocabulary and provides a framework for modeling conserved and species-specific gene-expression programs. **d**, Detailed architecture of Speciesformer.

Speciesformer adopts an encoder–decoder architecture tailored to single-cell modeling and controllable generation (Fig. 1a,d). The encoder is a deep transformer trained with self-supervision to capture macrogene-macrogene dependencies from multi-species expression profiles while integrating cell metadata and gene sequence priors. We developed several Speciesformer encoders of different size by using the same transformer encoder architecture (Supplementary Table 2). The decoder is a modality-flexible module that conditions on encoder representations and natural-language prompts to generate (Fig. 1a): (i) concise cell-state description, (ii) counterfactual expression profiles under specified perturbations, and (iii) pseudo-cells sampled from target states described in text (Supplementary table 3). This division of model architecture allows the encoder to learn universal biological structure, while the decoder translates those representations into task-specific outputs (Fig. 1d). Full details of Speciesformer architecture are provided in Methods and Supplementary Notes.

The encoder operates on an macrogene-level input and uses bidirectional self-attention to model regulatory co-variation and context. Pretraining is driven by masked reconstruction on the SpeciesCorpus, with late-fused metadata to disambiguate biological and technical heterogeneity, and macrogene sequence embeddings to inject evolutionary signal. Because every cell is expressed in the same macrogene frame, the model learns species-invariant correspondences while preserving species-specific residuals, yielding cell embeddings and gene embeddings suitable for downstream annotation, perturbation modeling, and regulatory analysis.

Built as a unified cross-modal generative module, the decoder operates on shared representations from the Speciesformer encoder together with task-specific biological conditions, enabling generation across three complementary cellular scenarios. For cell-state description, the decoder conditions on the encoded cell representation and generates a textual statement describing the biological state of the cell. In the reverse direction, for pseudo-cell generation, it conditions on a textual prompt that summaries the desired cell state, such as “CD14 cells derived from the blood tissue of an adult male human, generated using Smart-seq2 under healthy status,” and generates the corresponding gene-expression profile. For perturbation-conditioned generation, the decoder jointly incorporates the representation of the initial control cell state and a structured or textual description of the intervention to generate the full post-perturbation expression profile. These settings allow the same decoder framework to connect cellular states with biological semantics and to model transitions between cellular states under defined biological conditions.

### Speciesformer encoder provides biological cell and gene embeddings that generalize across diverse downstream tasks

To validate whether the learned cell and gene representations from Speciesformer encoder can capture shared biological structure across evolutionarily diverse species, we assessed their utility across multiple biological settings. At the cell level, we tested whether the learned representations preserved cell identity and generalized to held-out datasets. At the gene level, we examined whether the embeddings encoded biologically meaningful relationships among genes, including perturbation-associated functional patterns and gene interactions. We attached a task-specific prediction head for each downstream task and optimized the model using task-dependent fine-tuning protocols tailored to the corresponding pretraining setting (Methods).

We first evaluated cell type annotation on five public datasets of human tissues in two scenarios. We visualized the clustering results of Speciesformer cell embeddings for immune (Fig. 2b) and Pancreas (Noted as hPancreas, Fig. 1c) datasets, demonstrating well-segregated cell-type islands with tight within-type cohesion (Other three datasets in Supplementary Fig. 1a,c,e). For the alpha and beta cell types of hPancreas (Fig. 2c), Speciesformer can group cells of the same type into the same cluster, while the scGPT model divides them into three clusters, indicating the stronger class-consistent geometry of Speciesformer in the latent space. Across three intra-datasets, Speciesformer achieved high precision, with the Immune reference showing especially strong per-class performance (Fig. 2a,b, Supplementary Fig. 1b,d). On the inter-dataset hPancreas benchmark characterized by pronounced batch heterogeneity, Speciesformer maintained more than 0.90 accuracy for most cell types. Performance naturally dipped for rare categories (e.g., epsilon and mast cells), where accuracies were 0.60-0.70 due to the class imbalance and limited training signal. We further benchmarked the fine-tuned Speciesformer against scGPT(Cui, et al., 2024), Geneformer(Theodoris, et al., 2023), UCE(Rosen, et al., 2026), and CellFM(Zeng, et al., 2025) (Fig. 2a). Speciesformer significantly boosted the ability of cell type annotation compared to alternative methods in overall accuracy, achieved the highest macro-F1 on four of five datasets.

**Figure 2.**
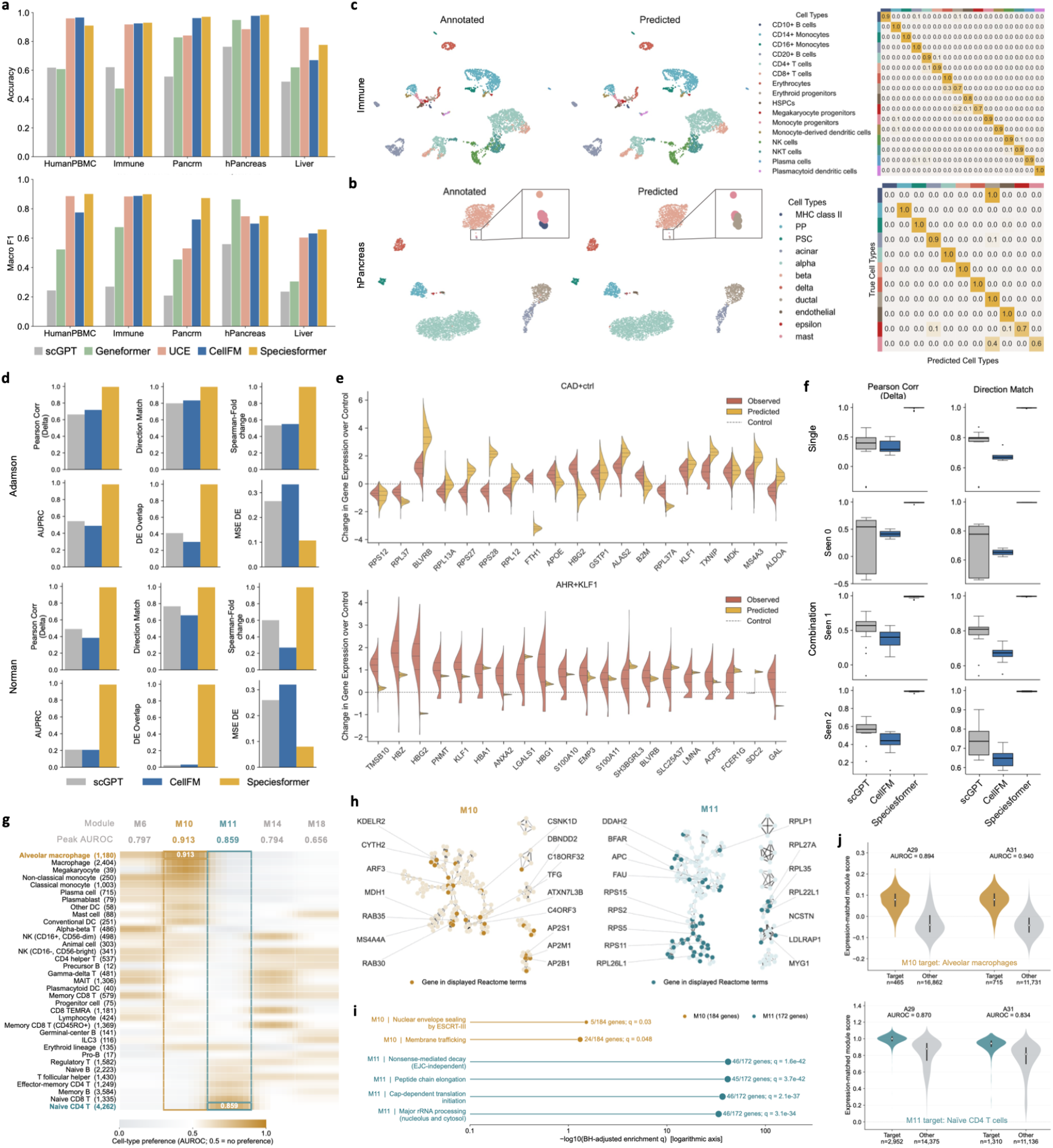
Speciesformer encoder provides biologically informative cell and gene embeddings across downstream tasks. Cell type annotation results on the Immune dataset (**a)**, and human pancreas dataset (**b**). UMAP visualization (left) comparing the ground-truth cell type labels and Speciesformer-predicted labels, colored by the cell types, and confusion matrix (right) for cell type annotation. **c**, Benchmarking of cell type annotation performance across five datasets. Accuracy and macro F1 score are reported for three intra-dataset settings and two inter-dataset settings. **d**, Genetic perturbation effect prediction performance on the Adamson *et al*. dataset (upper) and the Norman *et al*. dataset (down). **e**, Recovery of top 20 DE genes for representative perturbations of gene *CAD* in Adamson and the gene combination (AHR+KLF1) in Norman, comparing observed and predicted changes in post-perturbation expression relative to the control group. **f**, Two presentative evaluation metrics across four testing scenarios on Norman dataset. **g**, Discovering and characterizing gene modules on Immune dataset. Of 25 original embedding-derived modules, all five with at least one globally BH-significant Reactome pathway are displayed. **h**, Within-module gene similarity networks. An undirected edge is displayed when either gene ranks the other among its top three positive-cosine neighbors within the module. **i**, Reactome overrepresentation functional annotations with the actual 5,000-gene background. Point size encodes overlap count, and text gives overlapping genes/module size and q. **j**, Activity in measured expression by donor. Colored violins show the target cell type, while gray violins show all other cells within the same donor.

We next evaluated whether Speciesformer could extrapolate from an unperturbed cellular state to the counterfactual expression profile induced by a specified genetic perturbation. We fine-tuned the model on two widely used CRISPR Perturb-seq datasets, Adamson(Adamson, et al., 2016), comprising 86 single-gene perturbations and 68,603 cells, and Norman(Norman, et al., 2019), comprising 105 single-gene and 131 double-gene perturbations across 91,205 cells. Models were trained on a subset of perturbations and evaluated on held-out perturbations, thereby testing generalization to unseen interventions. Across all gene-perturbation pairs, it achieved global Pearson correlations of 0.98 on Adamson and 0.99 on Norman, indicating strong agreement between predicted and observed perturbed profiles (Fig. 2d, Supplementary Fig. 2a,b). We compared Speciesformer with scGPT and CellFM across six metrics, which quantify at two complementary levels: global agreement across all gene-perturbation pairs and per-perturbation fidelity on the top 50 differentially expressed (DE) genes. Across the two datasets, Speciesformer achieved the best performance across all six metrics. The advantage was particularly evident on the Norman dataset, where double-gene perturbations introduce combinatorial and potentially non-additive effects. In this setting, Speciesformer maintained highest correlation and directionality across four perturbation scenarios, whereas the baseline models showed substantially reduced performance (Fig. 2f). We further visualized two representative perturbation conditions, including a single-gene perturbation CAD in Adamson and a double-gene perturbation AHR+KLF1 in Norman. Fine-tuned Speciesformer recovered both the direction and magnitude of expression changes among the top 20 DE genes, whereas scGPT and CellFM produced predictions close to zero, suggesting conservative shrinkage toward the control state and limited ability to capture combinatorial perturbation effects (Fig. 2e, Supplementary Fig. 2c, d).

To validate whether the pretrained gene embeddings already encoded regulatory and functional relationships before task-specific fine-tuning, we extracted gene embeddings from Speciesformer on a human immune dataset and constructed a dataset-level gene network analysis to characterize gene-gene interactions. The derived gene embeddings grouped functionally related genes into 25 distinct modules (Supplementary Fig. 3a). We presented the AUROC scores of the top 5 significant modules of Reactome in 25 modules across all 35 types of cells. Based on the distribution of cell type activity of these modules, M10 and M11 respectively show preferences for alveolar macrophages and naive CD4 T cells, and their peak AUROC is the highest among these 5 modules (Fig. 2g, Supplementary Fig. 3b). The two modules exhibit distinct functional compositions, where M10 is enriched in membrane transport and the ESCRT-III-mediated nuclear membrane closure process, while M11 mainly involves translation, nonsense-mediated mRNA degradation, and rRNA processing (Fig. 2h,i, Supplementary Fig. 3d). In the measured gene expression, the average correlation within the modules of both was higher than that of randomly selected gene sets with matching expression means and variances (999 permutations, all with BH q = 0.0125; Supplementary Fig. 3c). Additionally, the cell type activity preferences of M10 and M11 were consistently in the same direction in the two donors, and the target cell type AUROC of the combined cells was 0.913 and 0.859 respectively (Fig. 2j).

### Speciesformer enables bidirectional generation between cell state and textual descriptions in human immune cells

We next validate whether these representations could support generative modeling of cell states. Recent multimodal single-cell models connect transcriptomic profiles with natural language, but bidirectional generation between cell state and text remains challenging. We therefore evaluated Speciesformer on two complementary tasks, generating cell type descriptions from expression profiles and generating pseudo-cell expression profiles from cell-state descriptions, on a human cross-tissue immune dataset(Conde, et al., 2022) comprising 29,790 cells and 34 cell types. We split the human immune dataset into training, validation and test sets at a ratio of 8:1:1 at both two scenarios.

For cell type description generation, cell expression profiles were encoded by the cross-species foundation encoder, and the resulting cell representation conditioned the generative decoder to produce a textual description of cell identity. To standardize cell type description, we referenced the cell types into ontology terms derived from “The Cell Ontology”(Bard, et al., 2005). On 2,978 test cells, the model showed strong cell-state understanding and language generation performance. Speciesformer generated descriptions that were broadly consistent with the annotated cell identities and captured key biological attributes of the input cells (Fig. 3a). After extracting predicted cell type labels from the generated descriptions using rule-based matching, Speciesformer achieved a 73.4% exact label match across the 34 annotated cell types, with 2,185 test cells correctly mapped to their ground-truth cell type (Fig. 3a). Quantitatively, Speciesformer outperformed GPT-2(Alec Radford, 2019), C2S(Levine, et al., 2024) and scMMGPT(Jiaqi Yang, 2026) with 41.95% improvements averagely over the second-best method (Fig. 3b, Supplementary Table 6).

**Figure 3.**
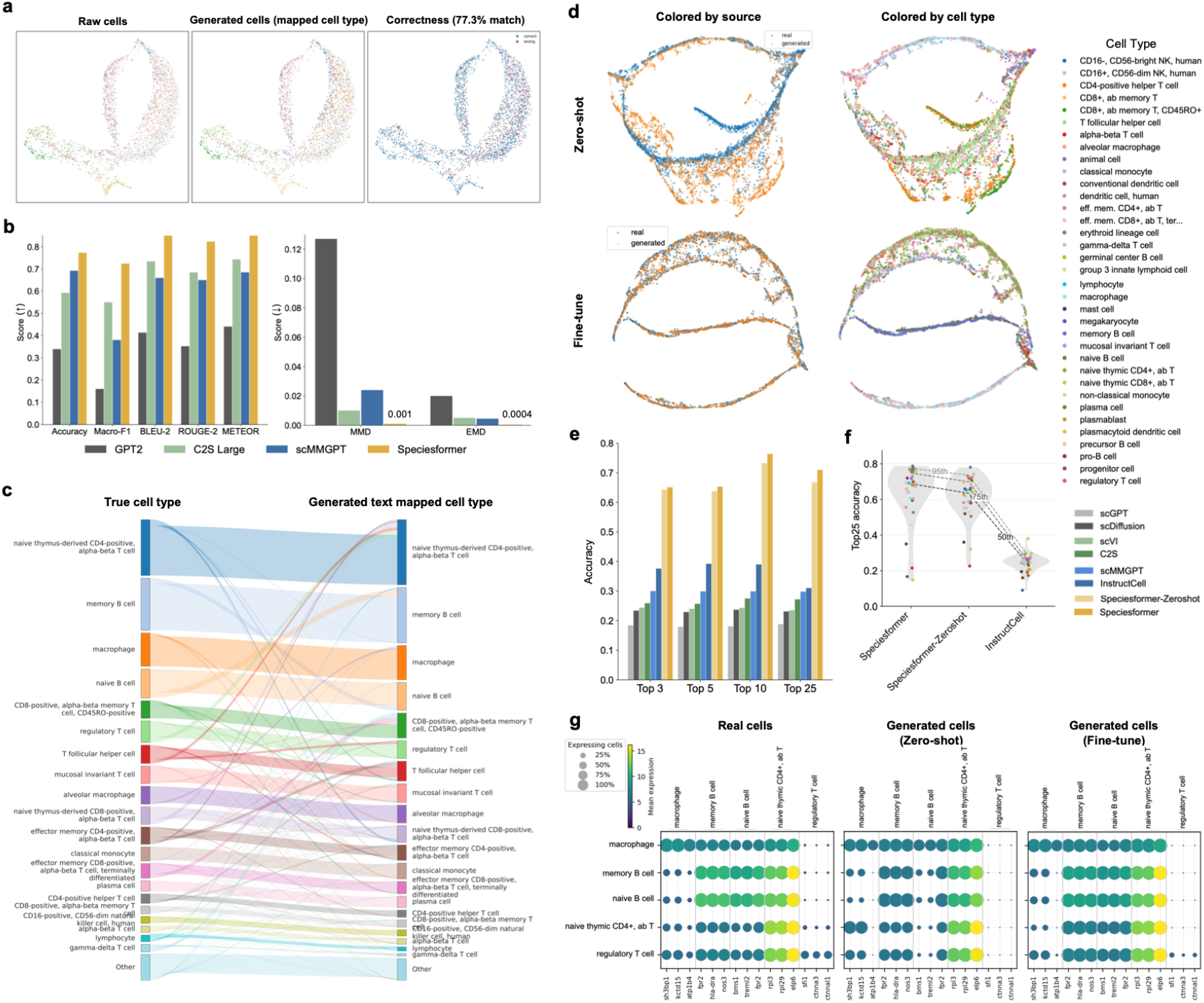
Speciesformer enables bidirectional generation between cell state and textual descriptions in human immune cells on the human cross-tissue immune dataset. (**a-c**) Speciesformer generates cell-type descriptions from single-cell expression profiles: **a**, UMAP visualizations compare ground-truth cell types and generated cell-type descriptions mapped back to cell-type labels. **b**, Benchmarking of cell-type description generation against three single-cell multimodal large models. (Left) Evaluation on five metrics, where higher values indicate better performance. (Right) Evaluation on two distance metrics, where lower values indicate better agreement between generated and reference textual distributions. **c**, Sankey diagram linking ground-truth cell types to the cell types mapped from generated cell descriptions. The flow widths summarize the correspondence between annotated cell identities and the semantic cell-type assignments produced by Speciesformer. (**d-g**) Speciesformer generates pseudo-cell expression profiles from textual descriptions of cell states: **d**, UMAP visualizations show pseudo-cells expression generated by the zero-shot model and the fine-tuned model from cell-state descriptions. Cells are colored by source (left) and by cell type (right). **e**, Cell-type transfer evaluation between real cells and generated pseudo-cells. **f**, Per-cell-type F1 scores distributions across zero-shot and fine-tuned Speciesformer, and second-best method. **g**, Dot plots of gene expression patterns for real cells, zero-shot and fine-tuned generated pseudo-cells. For each cell type with more than 150 cells, the top three significant genes were identified using Welch’s t-test and displayed along the x-axis.

We further examined where correct and incorrect predictions occurred in the expression and text representation spaces. At the cell type level, Speciesformer performed well on major immune populations, including plasma cells, macrophages, memory B cells, classical monocytes and naive CD4 T cells (Fig. 3c). The main sources of confusion occurred among B cell subtypes and among fine-grained T cell populations, including CD4 and CD8 T cells, regulatory T cells, T follicular helper cells, and effector or memory T cell subsets (Supplementary Fig. 4a). In the text embedding space, generated descriptions and target descriptions were generally located within nearby semantic clusters, indicating that the generated text preserved the overall semantic structure of the reference descriptions (Supplementary Fig. 4b).

To validate that Speciesformer enables conditional pseudo-cell generation from cell states descriptions, we employed Speciesformer under both zero-shot and fine-tuned settings to generate pseudo-cell expressions based on cell state descriptions of immune test set. UMAP visualization showed that zero-shot generated cells broadly recovered the major manifold structure of real immune cells, although local source-specific shifts and partially separated branches remained. Fine-tuning substantially improved the mixing of generated and real cells, yielding more consistent global trajectories and cell type-associated local structure (Fig. 3d). Consistently, expression-level comparisons showed a more dispersed zero-shot prediction distribution, particularly in highly expressed regions, whereas fine-tuning concentrated the high-density region closer to the diagonal and reduced prediction dispersion (Supplementary Fig. 5). We further benchmarked two Speciesformer settings against leading single-cell generative models: scVI (Lopez, et al., 2018), scDiffusion (Luo, et al., 2024), scGPT (Cui, et al., 2024), C2S (Levine, et al., 2024), scMMGPT (Jiaqi Yang, 2026), InstructCell (Fang, 2025) (Fig. 3e). To conduct a fair benchmarking, we trained a naïve k-nearest neighbors cell type classifier by using the test set, and predicted cell type from the cell expression generated by each model. Quantitatively, both zero-shot and fine-tuned Speciesformer substantially outperformed the baseline models with improvement of 62.5% to 95.9% than second-best model InstructCell across multiple evaluation metrics (Fig. 3e,f). Fine-tuning further improved all recorded test top-k accuracy metrics over the zero-shot setting. Specifically, top-3 and top-5 accuracy increased by approximately 0.8% to 1.6%, while top-10 to top-25 accuracy improved by approximately 3.1% to 4.2% (Fig. 3e,f). We further evaluated whether generated pseudo-cells recovered cell type-specific marker expression patterns. We filtered five cell types with more than 150 cells, and identified the top3 marker genes from the real cells and compared their expression patterns with those in zero-shot and fine-tuned generated cells for each cell type. Both settings reproduced the major marker-gene signatures, while fine-tuning improved the agreement with real cells in both mean expression levels and the fraction of cells expressing each marker, particularly for B cell- and T cell-associated genes (Fig. 3g).

### Speciesformer generates perturbation-conditioned cellular responses

A central objective of virtual cell modeling is to predict how cellular states change after perturbation, particularly in cellular contexts that have not been experimentally profiled. Because perturbation responses are highly context dependent, a useful model should generalize beyond observed cell types, cell lines or biological settings rather than simply interpolate within familiar contexts. Computational models have been developed to predict perturbation-induced expression changes from single-cell perturbation atlases. While perturbation prediction has traditionally been formulated as a supervised mapping(Lotfollahi, et al., 2019)(Bunne, et al., 2023)(Roohani, et al., 2024)(Ji, et al., 2021), generative foundation models further provide the ability to simulate perturbation-conditioned cell states in a context-aware manner(Theodoris, et al., 2023)(Cui, et al., 2024)(Hao, et al., 2024)(Pearce, et al., 2026)(Adduri, et al., 2025)(Klein, et al., 2025)(Dong, et al., 2026). However, robust generalization remains challenging when perturbation effects must be transferred across cellular contexts (Ahlmann-Eltze, et al., 2025), due to the substantial biological and technical heterogeneity(Wei, et al., 2026). These limitations motivate a formulation that treats perturbation response prediction not only as regression from one expression profile to another, but as conditional generation over cellular state space. We therefore formulate perturbation response prediction in Speciesformer as conditional perturbed-cell generation, where the model generates the post-perturbation expression state from a control-cell representation together with a textual description of the perturbation condition. This formulation enables us to directly evaluate whether Speciesformer can transfer perturbation dynamics across cellular contexts within a unified generative framework.

To validate the universal capability of Speciesformer for perturbation-condition cellular response prediction, we benchmarked Speciesformer across multiple large-scale perturbation datasets(Zhang, et al., 2025)(Srivatsan, et al., 2020)(Biosciences, 2023), covering settings from cross-context generalization of perturbation responses to transfer of perturbation dynamics in entirely new contexts. We compared Speciesformer with a broad set of baselines, including CPA(Lotfollahi, et al., 2023), scVI(Lopez, et al., 2018), context-mean baselin, and two leading perturbation prediction models, STATE(Adduri, et al., 2025) and CellFlow(Klein, et al., 2025). All methods were trained on the same data splits using their official implementations and evaluated under a unified Cell-Eval protocol(Adduri, et al., 2025), which provides a multi-level evaluation strategy that assessed full-expression reconstruction, perturbation-induced expression shifts, differentially expressed gene recovery and preservation of perturbation-specific effect sizes. We reported multiple complementary metrics, including Pearson correlation (delta), perturbation discrimination score (L2), differential expression overlap accuracy, differential expression precision, spearman correlation (Fold change), AUROC and AUPRC, to assess both global expression reconstruction and perturbation-specific response recovery.

We first investigated whether Speciesformer could generalize perturbation responses to new cellular contexts. We evaluated this setting on Tahoe-100M(Zhang, et al., 2025), a large-scale chemical perturbation dataset containing responses from 50 diverse cancer cell lines treated under 1,138 conditions involving 380 distinct drug perturbations. We employed the same evaluation strategy as used in STATE (Fig. 4a), measuring whether limited observations from a new cellular context are sufficient for a model to adapt its perturbation predictions beyond fully observed training contexts. We first assessed whether models could recover perturbation-induced expression changes. Speciesformer achieved the highest scores on two correlation-based metrics, with PCC(Delta) of 0.452 and LFCSPCC of 0.454, outperforming the second-best model, mean-based baseline, which reached 0.211 and 0.197 (Fig. 4b, Supplementary Fig. 8a). We next evaluated whether the predicted responses were sufficiently specific to distinguish different perturbation effects. Speciesformer achieved the highest PDS(L2), with a score of 0.798, compared with 0.564 for the strongest baseline. Consistent with this result, Speciesformer also obtained the highest AUPRC and AUROC, reaching 0.597 and 0.701, whereas the second-bast baseline scores were 0.52 and 0.514 (Fig. 4b, Supplementary Fig. 6). We further found that the performance of Speciesformer was consistently above mean-based baseline across cell lines, development phase, drug perturbations, and doses (Fig. 4c, Supplementary Fig. 8b-d). Moreover, Speciesformer’s AUPRC and AUROC are 36.2% and 25.7% higher than the second-best approach scVI on the PANC-1 cell line, respectively (Fig. 4d). We further examined recovery of differentially expressed genes, which evaluates whether models identify genes most strongly affected by the intervention. Speciesformer was particularly effective at recovering the highest-confidence perturbation-response genes. It achieved the best DE-gene overlap accuracy and precision at top 50, 100 and 200, showing that it more accurately prioritized the most strongly perturbed and biologically informative response genes (Supplementary Fig. 7). Although some baselines performed better at broader cutoffs such as top 500 and all DE genes, these larger sets may include weaker-effect or less perturbation-specific genes (Fig. 4b, Supplementary Fig. 6,7). Overall, these results suggest that no model dominated every individual metric, but Speciesformer provided the strongest balance across the evaluation levels most directly associated with biologically meaningful perturbation generation.

**Figure 4.**
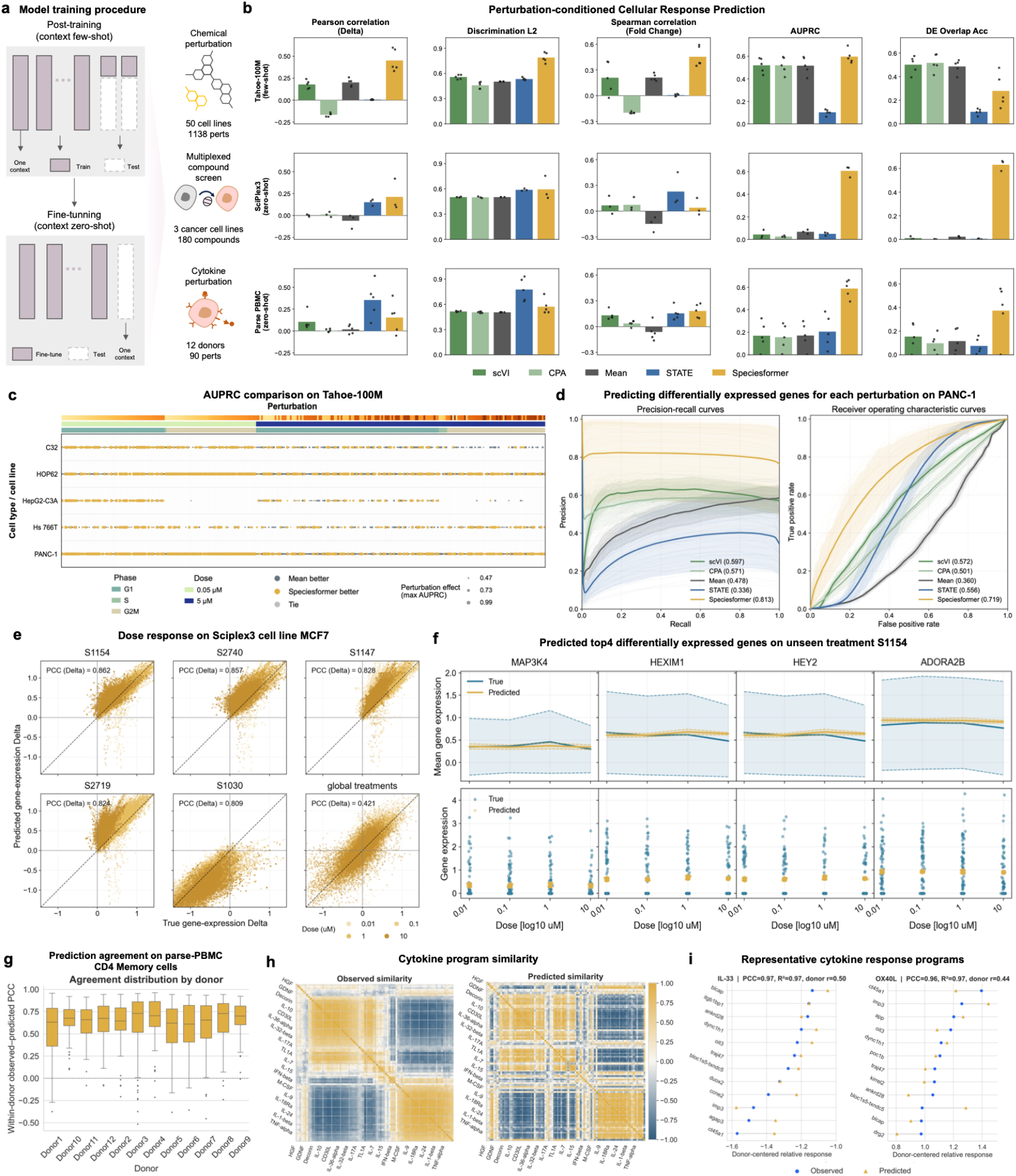
Speciesformer outperforms existing baselines in predicting perturbation effect across context generalization tasks. **a**, Data split on model training and evaluation procedure at two context generalization setting. **b**, Perturbation-conditioned cellular response prediction performance across three perturbation datasets. Bars represent the mean metric value across evaluated cell type/cell line groups, and overlaid dots show individual group-level aggregate values. **c**, Dot plot comparing Speciesformer with the Mean baseline across perturbations on the Tahoe-100M dataset. Dot color indicates which method performs better, while dot size represents AUPRC value. Perturbations are organized by dose and cell-cycle phase. **d**, Precision-recall (PR; left) and receiver operating characteristic (ROC; right) curves on the PANC-1 cell line across perturbation-dose conditions shared by all methods. **e**, Dose-dependent response generation in sciPlex3 MCF7 cells for the top5 treatments, and the global aggregating treatments. **f**, Dose-dependent expression responses to the unseen drug S1154. True and Speciesformer-predicted expression profiles are shown across four doses for representative differentially expressed genes. **g**, Predictive consistency of cytokine response programs across biological donors. Each box plot summarizes the observed-predicted program PCC of 90 cytokines within a donor. **h**, Similarities among different cytokine relative response programs and their predictive recovery. **i**, Two representative donor-centered relative response programs (IL-33, and OX40L) on 12 highly contributing genes.

We next assessed whether Speciesformer could transfer perturbation dynamics to entirely new cellular contexts. This setting, which we refer to as zero-shot perturbation prediction across contexts, is closer to the central goal of virtual cell modeling. We therefore asked whether post-training on Tahoe-100M could provide transferable perturbation-response knowledge that improves prediction in previously unperturbed cellular contexts. To test this capability, we employed the Speciesformer model post-trained on Tahoe-100M and fine-tuned it on two smaller perturbation datasets: SciPlex3 (Srivatsan, et al., 2020), a multiplexed compound screen dataset, and Parse-PBMC (Biosciences, 2023), a cytokine perturbation dataset. For each dataset, we held out one cell context at a time during training and evaluated prediction performance in the held-out context.

We first evaluated zero-shot context-level perturbation prediction on SciPlex3. Speciesformer showed the strongest performance in recovering differentially expressed genes (Fig. 4b, Supplementary Fig. 6,7). It achieved DEOAcc of 0.631 and DEPrec of 0.607, whereas all baseline models remained substantially lower. This result indicates that Speciesformer more effectively recovered the genes most affected by drug perturbations in held-out cellular contexts. At the level of perturbation-induced expression shifts, Speciesformer achieved the highest PCC(Delta), reaching 0.213, indicating stronger agreement between predicted and observed transcriptional changes than the compared methods (Fig. 4b, Supplementary Fig. 9a). This advantage was also maintained across dose-stratified evaluations, where Speciesformer consistently recovered perturbation-response patterns across different treatment doses (Supplementary Fig. 9b). For perturbation-level discrimination, Speciesformer achieved the highest AUPRC, with a score of 0.608, and the highest PDS(L2), with a score of 0.595. These results indicate stronger recovery of perturbation-specific response structure and better discrimination of drug-induced states in held-out cellular contexts. To investigate the ability of modeling continuous dose-response relationships, we examined the post-perturbation predictions from Speciesformer on top 5 and global unseen treatments for MCF7 cells. The predicted changes in gene expression by the model were highly consistent with the actual changes in gene expression (Fig. 4e). On an unseen treatment S1154, Speciesformer accurately captured the changing trends of different differentially expressed genes as the drug concentration increased, and simultaneously recovered the central range of the perturbed state of real cells (Fig. 4f).

We further evaluated zero-shot context-level perturbation prediction on Parse-PBMC, a cytokine perturbation dataset. Compared with SciPlex3, this task examines whether the model can generalize perturbation dynamics to an immune-cell setting and a non-chemical perturbation modality. Speciesformer showed the strongest recovery of perturbation-responsive genes, achieving DEOAcc of 0.374 and DEPrec of 0.373, with all baseline methods performing lower on both metrics (Fig. 4b, Supplementary Fig. 6,7). Beyond identifying responsive genes, Speciesformer also preserved perturbation-specific effect sizes, obtaining the highest LFCSPCC of 0.183, while ranking second on PCC(Delta) with a score of 0.154. At the perturbation-discrimination level, Speciesformer achieved the highest AUPRC of 0.588 and ranked second on AUROC and PDS(L2), with scores of 0.536 and 0.572, respectively (Fig. 4b, Supplementary Fig. 6). We further validated whether the cytokine relative program predicted by the model on CD4 memory cells could be consistently reproduced across different donors. Speciesformer showed good directional consistency in most donor-cytokine combinations, and there was no overall failure at the donor level, with PCC values on all donors higher than 0.61 (Fig. 4g, Supplementary Fig. 10a,b). Moreover, the model has successfully restored the global organizational structure of the cytokine-specific relative response program in CD4 Memory (Fig. 4h, Supplementary Fig. 10c). Particularly, for the two cytokines IL-33 and OX40L, which show a very strong inverse relationship in the real data, Speciesformer has almost completely restored this relationship (Fig. 4i, Supplementary Fig. 10c).

Together, these analyses indicate that Speciesformer can serve as a foundation for virtual perturbation modeling and cross-context prediction of cellular responses, while highlighting the need for continued evaluation across broader perturbation modalities, datasets and biological contexts.

## Conclusions

Speciesformer represents a cross-species generative virtual cell model designed to learn cellular states and state transitions while separating transferable biological programs from context-specific variation. By constructing SpeciesCorpus and mapping genes from multiple species into a shared macrogene vocabulary, Speciesformer establishes a common modeling space for conserved and species-dependent transcriptional programs. Its encoder learns biologically informative cell and gene representations that generalize across multiple forms of representation probing. Speciesformer achieved strong cell type annotation performance in both intra- and inter-dataset settings, recovered genetic perturbation-induced expression changes in Adamson and Norman, and identified gene modules with interpretable functional enrichment, indicating that the learned representations preserve biological structure at both cell and gene levels.

Building on this shared representation space, Speciesformer extends cross-species foundation modeling from representation learning toward generative modeling of cellular states. On the human immune dataset, the unified decoder supported bidirectional generation between transcriptomic states and biological text descriptions. Expression-to-text generation recovered cell identities consistent with reference annotations and performed strongly across semantic and distributional evaluations, whereas text-to-expression generation produced pseudo-cells that preserved cell type structure and marker-gene programs in a zero-shot setting, with fine-tuning further improving agreement with observed cells. Speciesformer further models perturbation-induced state transitions through conditional perturbed-cell generation. Across multiple perturbation datasets and evaluation settings, the model recovered response-associated genes, perturbation-induced expression changes and perturbation-specific response structure, while maintaining the ability to generalize across previously unobserved cellular contexts. These results extend the role of the shared cellular state space from describing observed cells to predicting how those states change under intervention.

Collectively, these findings position Speciesformer as a step toward a cross-species generative virtual cell framework. Rather than treating comparative analysis, biological representation probing, cell-state generation and perturbation prediction as disconnected problems, Speciesformer integrates them around a common objective of modeling cellular states and their context-dependent transitions. This formulation provides a basis for studying which cellular programs are broadly transferable across biological systems and which responses remain dependent on species, cell identity or experimental context.

Several directions remain important for future work. Large-scale pretraining benefits from increasingly heterogeneous single-cell collections, but batch effects and dataset-specific biases can still confound the biological variation that foundation models aim to capture, and handling these factors in a unified manner remains a shared limitation of current foundation models. Furthermore, incorporating spatial transcriptomics into the modeling framework allows virtual cell models to move beyond isolated expression profiles and account for tissue architecture, local neighborhoods and spatially constrained cell-cell interactions. Integrating spatial context with cross-species and generative modeling could ultimately allow virtual cell models to represent not only what a cell is and how its state changes, but also where those states arise and how tissue organization shapes cellular behavior.

## Methods

### Multi-Species Single-cell corpus 131M

#### Data curation and preprocessing

We assembled SpeciesCorpus, a large-scale multi-species scRNA-seq corpus for foundation model pretraining and evaluation, by integrating publicly available datasets from CELLxGENE Census and GEO across multiple species, tissues and sequencing protocols. Datasets were included when cell-by-gene count matrices were available and gene identifiers could be mapped to authoritative Ensembl or NCBI annotations, while associated metadata describing species, tissue, sex, developmental stage, disease state and library preparation were retained when available. To ensure consistent preprocessing across data sources, we applied source-agnostic quality control and restricted the analysis to Ensembl-annotated protein-coding genes. Cells with fewer than 200 detected protein-coding genes were excluded, and raw counts were subsequently library-size normalized and log1p transformed. Detailed dataset acquisition, inclusion criteria, metadata harmonization and quality-control procedures are provided in the Supplementary Notes.

#### macrogene construction strategy

To establish a shared gene space across multiple species, we constructed evolution-informed macrogenes rather than relying on pairwise one-to-one ortholog mapping. We employed kernel archetypal analysis(Cutler and Breiman, 1994) to derive cross-species macrogenes in a protein embedding space. Protein representations were obtained for genes from all species using the pretrained ESM2 protein language model(Lin, et al., 2023), providing a common functional embedding basis across lineages. We then organized macrogene construction according to the phylogenetic relationships among species, progressively aggregating homologous gene structure from lower to higher taxonomic levels through an inverse phylogenetic-tree strategy. This procedure yielded 862 macrogenes shared across the 11 species, and the learned gene-to-macrogene mappings were used to project species-specific expression matrices into a unified cell-by-macrogene space for pretraining and downstream analyses. Detailed procedures for protein embedding aggregation, neighborhood construction, archetypal decomposition and hierarchical macrogene extraction are provided in the Supplementary Notes.

### Speciesformer Architecture and pretraining

#### A foundation model as an Encoder

We design inputs to learn cell and gene-level representations while remaining robust across species. To unify homology across species, we first map genes to macrogenes (orthology-aware groups). The corpus uses a single macrogene vocabulary of size *M*. For every cell, we construct an *M*-length sparse count vector and aligned token list. That is, single-cell sequencing data across species are processed into a cell-by-macrogene matrix, *X* ∈ ℝ^*N*×*M*^. Because every cell indexes the same vocabulary, no padding is required for macrogene tokens, macrogene expression values, or macrogene sequence embeddings. The input to Speciesformer consists of four main components: (1) macrogene tokens, (2) macrogene expression values, (3) macrogene sequencing embeddings, and (4) cell metadata.

#### Macrogene tokens

As we assume that the gene expression profile of a cell is like a sentence, composed of “words” like genes. In our framework, different cellular sentences are composed of fixed macrogene tokens, only the expression levels of each macrogene differ. Each macrogene mg has been assigned to a unique integer identifier id. The approach offers stable, context-agnostic attributes and keys/queries within a cell, analogous to the “gene tokens” used by scGPT and the rank-encoded gene tokens of Geneformer. In addition, we incorporate special tokens <CLS> in the vocabulary for aggregating all macrogenes into a cell representation. The input macrogene tokens of each cell *i* are hence represented by a vector 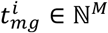, M is the number of macrogenes in this study.

#### Macrogene expression embeddings

The unified preprocessed gene expression matrix needs to be transformed into macrogene level before it can be encoded. Let the set of species be *S*, and the species be *s* ∈ *S*. For the expression value of the *j*-th “intraspecific gene” in the *i*-th cell of this species, represented as 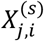, we used the previously obtained gene-to-macrogene mapping to perform cross-species alignment of gene expression in different species. Let the mapping from genes within a species *s* to macrogenes be denoted as *ϕ*_*s*_: *j* ⟼ *mg* ∈ {1, 2, …, *M*}, *G*^s^(*mg*) = {*j*: *ϕ*_*s*_(*j*) = *mg*}. Then, the member genes of the same macrogenes in species *s* are aggregated into a macrogene group:

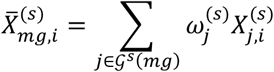

Where 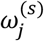 is the weight from gene *j* to macrogene *mg*. Because we use a uniform metagenomic vocabulary size *M*, for any cell *i* and species *s*, 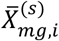 are fixed-length vectors, therefore padding is unnecessary. For each cell, the normalized value of macrogene mg is transformed into a value embedding by a two-linear-layer MLP with GELU for continuous projection, enabling smooth sensitivity to dynamic range:

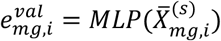

We then form a macrogene-slot embedding by additive fusion:

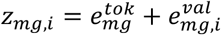

The use of macrogene tokens plus expression signals remains compatible with masked-value or regression objectives.

#### Macrogene sequencing embeddings

To expose cis-sequence context, each gene carries a combined embedding 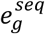 precomputed and aggregated from its representative transcript sequence using several pre-trained large language models (Supplementary Notes). Gene-level embeddings were lifted to the macrogene vocabulary per species using the precomputed mapping *ϕ*_*s*_. When multiple genes mapped to the same macrogene in a species, their embeddings were combined by a weighted average using the learned weights. This produced one DNA-sequence, one protein-sequence, and one text-description vector per macrogene.

To form a single macrogene sequence embedding, the three modality-specific vectors were first independently normalized with LayerNorm, then concatenated along the feature axis and projected to a fixed dimension *d* through a linear layer. A final LayerNorm was applied to the projected vector to stabilize scale across macrogenes and species. Formally, letting 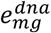, 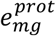, and 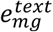 denote the macrogene-level embeddings for DNA, protein, and description respectively,

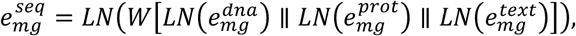

Where [·∥·] denotes concatenation, *W* is a learnable projection to ℝ^*d*^, and *LN* is LayerNorm. The resulting 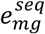 is cached and reused all cells expression macrogene *mg*. At model input, the fused macrogene sequence embedding 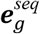 is added to the corresponding macrogene token and value embeddings, yielding the final slot embeddings for macrogene *mg*.

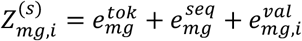

### Cell metadata tokens

In addition to macrogene-level tokens, we introduce cell metadata tokens to encode contextual factors that modulate cellular state and measurement. These tokens capture species, assay, tissue, sex, developmental stage, disease state, and sample-level technical signals, enabling the model to condition on biological and technical heterogeneity. Metadata are harvested from dataset-provided annotations and standardized to controlled vocabularies. Specifically, species are mapped to the Taxon ontology (NCBI Taxonomy IDs), tissues and organs to UBERON ontology, and disease labels to MONDO ontology. We utilized the accepted assay documents provided by CZI CELLxGENE census to normalize assay chemistry labels. For all species, we only employed three gender labels: female, male, unknown, and five development stage labels: “Fetal”, “Juvenile”, “Neonatal”, “Adulthood”, and “unknown”. If a field is missing or irreconcilable, its value is set to “unknown”. Each metadata attribute *a* is associated with a discrete vocabulary *V*_*a*_ that includes an unknown token. Values are embedded via independent learnable tables:

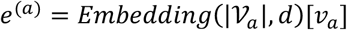

where *v*_*a*_ ∈ *V*_*a*_ is the normalized category and *d* is the model width defined for cell metadata tokens. Unknown categories are treated as first-class tokens and are included during training to make the model robust to incomplete or partially harmonized metadata. Let {*e*^(*a*)^}_*a*∈*A*_ denote the embeddings for available metadata attributes (species, assay, tissue, sex, developmental stage, disease, technical signals). We aggregate metadata into a single vector by addition:

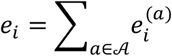

### Speciesformer encoder

#### Model architecture

We adopt a self-attention transformer encoder to model macrogene–level dependencies from inputs embeddings. The network comprises *T* encoder blocks, each with multi-head self-attention followed by a feed forward neural network layer. Inputs are fixed length to 862 macrogene slots plus a single <CLS> token that aggregates cell-level context. Regularization includes 0.1 hidden dropout and 0.1 attention dropout. Weight matrices are initialized with standard deviation 0.02, and layer normalization use epsilon of 1×10^−12^.

The final encoder outputs a *D*-dimensional representation per macrogene and a *D*-dimensional representation at <CLS>. We regard this special token as a learned pooling operation within transformer blocks. Thus, we use the representation of <CLS> as cell representation as it aggregates the learned macrogene-level representations. To encode sample context without perturbing the fixed macrogene layout, cell metadata vectors are fused at the prediction head: per-macrogene hidden states and the <CLS> embedding are concatenated with the metadata vector only for decoding. Metadata therefore conditions predictions post-hoc, improving context awareness while preserving a strictly fixed macrogene frame and the transformer’s attention topology.

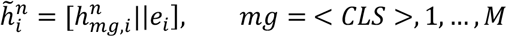

### Speciesformer encoder pretraining

Pretraining for Speciesformer encoder follows a masked-learning scheme tailored to macrogene profiles. For each cell, we mask 15% of macrogenes by withholding their tokens and expression values. The model reconstructs masked expressions from the unmasked context via two complementary heads: (1) Macrogene-slot head applying to each masked macrogene’s hidden state, regressing its expression. (2) Cell-context head applying to the <CLS> embedding, projecting to a full-length vector and regressing the masked entries. Both heads optimize mean squared error (MSE) on the chosen value space, and the total loss is the sum:

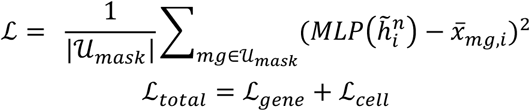

Where 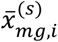 denotes the truth macrogene expression value under masking and *U*_*mask*_ is the set of masked positions of macrogenes. This design encourages slot-wise precision while leveraging global cell context, and late-fused metadata conditions both decoders on species, tissue, assay, and other factors without altering macrogene indexing.

### Speciesformer encoder fine-tuning

Because most downstream objectives differ from the masked-reconstruction pretraining objective, we adopt task-specific fine-tuning or post-training with lightweight decoders while adapting the pretrained encoder. Unless stated otherwise, we initialize all fine-tuning models from the pretrained Speciesformer encoder checkpoint, replace the pretraining head with a task-specific decoder, and use late fusion of metadata exclusively at the decoder.

#### Cell type annotation

Accurate cell-type annotation remains a central bottleneck for translational single-cell analysis, particularly in cross-species settings where marker usage and granularity differ. We frame fine-tuning annotation as a supervised classification problem over the <CLS> representation augmented with contextual metadata to account for diverse variability. We attach a two-layer MLP classifier to the concatenation of the <CLS> embedding and the metadata vector. The objective of this task is to optimize the classification loss using the cross-entropy between the predicted cell type probability and the true label. Because the objective differs substantially from masked reconstruction, all encoder layers are unfrozen, while classifier weights are randomly initialized.

#### Perturbation response prediction

Predicting cellular responses to genetic or pharmacological perturbations is key to hypothesis generation and therapy design. Unlike masked value imputation, perturbation modeling requires extrapolating from a control cell state to a counterfactual perturbed state conditioned on a perturbation specification. The pretrained encoder processes the control macrogene profile. A perturbation token encodes the targeted macrogene(s) as a sparse vector. For this task, we construct a perturbation-specific regression head outputting post-perturbation macrogene profiles. Training minimizes MSE on predicted profiles against observed perturbed profiles. Given partial alignment with pretraining, we freeze embeddings and the first *K* transformer blocks, updating only the last *T* − *K* blocks and the perturbation head. This preserves base representations while allowing higher-layer adaptation to causal conditioning.

#### Gene network analysis

For the gene network analysis task, we extract cell-level macrogene embeddings and convert them into dataset-level gene vectors by averaging across cells. A KNN graph (cosine similarity) over genes defines the similarity network. We apply Leiden clustering and retain clusters with ≥5 genes as gene programs. Cluster profiles are summarized by average expression and enrichment of known pathways.

### A Unified Generative Decoder

Bridging from representation to generation, the encoder results establish that Speciesformer learns cell and gene-level structure that is linearly decodable, robust across datasets, and richly organized in latent space. We leverage these representations with a generative decoder that translates biology-aware embeddings into controllable virtual transcriptomes. Speciesformer couples a macrogene-aligned, multi-species encoder with a prompt-conditioned generative decoder to unify three capabilities: (i) cell semantic modeling, where the decoder renders concise textual descriptions grounded in the cell embedding, or (ii) text-prompt to pseudo-cell synthesis, which generates full macrogene expression vectors from natural-language prompts specifying target states, (iii) conditioned-perturbation response generation, which maps a control profile plus a textual perturbation specification to the post-perturbation expression profile. Throughout, all outputs are emitted in the same orthology-aware macrogene space, enabling direct, quantitative comparisons across species, tissues, and assays-functionality that, to our knowledge, has not been demonstrated jointly by prior virtual-transcriptome models.

#### Architectural overview

Concretely, we first extract the cell embeddings *h*_*cell*_ and per-macrogene slots *H* ∈ ℝ^*M×d*^ from the Speciesformer encoder, and obtain the text embeddings *t* from a pretrained Llama3.1-8B model(Patterson, et al., 2022). Then, *h*_*cell*_ and *t* are mapped into a shared latent via two small projectors and fuse them with gated cross-attention in the decoder head. We froze backbones and employed light cross-modal adapters to let most trainable parameters reside in the projectors and task heads, enabling efficient adaptation without changing the macrogene layout.

#### Cross-modal projectors

For each cell, the encoder yields a global cell embedding ***h***_*cell*_ ∈ ℝ^d^ and per-macrogene hidden states *H* = [**h**_1_; …; **h**_*M*_] ∈ ℝ^*M×d*^. A text prompt is tokenized by the LLM to a sequence 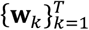 with hidden size *d*_*t*_. We form a pooled text representation 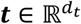 via the LLM’s recommended pooling. We map ***h***_*cell*_ and, when needed, each ***h***_*Mg*_ into the shared latent by a two-layer MLP with residual scaling and LayerNorm:

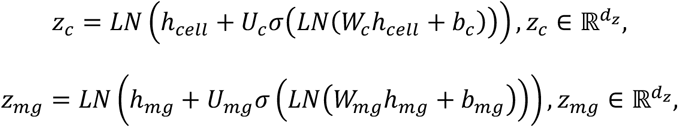

where *W*_·_, *U*_·_ are learnable matrices, *b*_·_ biases, *σ* is GELU, and *LN* is pre-activation LayerNorm. We employed dropout after the hidden layer to improve stability under mixed-domain training. The pooled text embedding is transformed analogously:

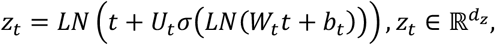

For tasks requiring token-level conditioning, we also project each LLM token ***w***_*k*_ to 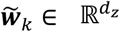 with shared weights (*W*_1,_ *U*_*t*_), enabling sequence-aware fusion.

Projected latent is fused by a light multi-head cross-attention with learnable gates that regulate modality influence:

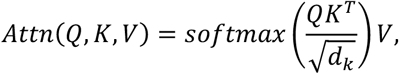

For cell-type description generation, queries arise from the text decoder states, keys/values from 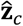 (or 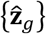 for gene-aware captions). For pseudo-cell and counterfactual generations, queries are the macrogene states {**h**_*g*_} or their projections 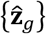; keys/values are 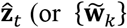 for token-wise control). A scalar gate *α* ∈ [0,1] per head and per block, obtained from a sigmoid of a learned parameter *θ*, blends attended and residual features:

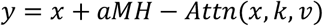

followed by a position-wise MLP and residual LN. Gates prevent over-conditioning and improve robustness when prompts are underspecified. For counterfactual generation, cross-attention outputs parameterize a *delta* head that predicts 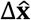. We provide the head both the control macrogene features (keys/values from *H* or 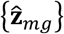) and the projected text prompt 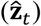, enabling interactions between what to change (text) and what exists (control).

To reduce modality anisotropy, we apply *LN* in projectors and include a contrastive alignment between **z**_*c*_ and the pooled text **z**_1_ for paired (cell, caption) data:

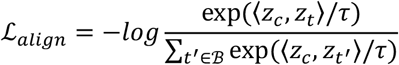

with temperature *τ* learned and in-batch negatives *B*. For pseudo-cell generation, a cycle-consistency term encodes 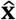 through the frozen encoder and pulls the resulting cell latent toward **z**_*t*_.

### Cell-type description generation

For this task, we aim to generate concise textual descriptions of cellular identity from learned cell embeddings. The decoder operates on the encoder-derived cell embedding, projected into the shared latent **z**_*c*_ = *p*_*c*→*z*_(**h**_cell_). A metadata vector *m* is incorporated by FiLM-style modulation of **z**_*c*_ yielding 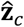.

Llama3.1-8B model is considered as the language head that conditions the token stream of a frozen LLM on 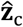 via gated multi-head cross-attention. Optionally, per-macrogene projections {**z**_M*g*_} augment 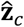 to enable gene-aware attributions. We minimize sequence cross-entropy under teacher forcing,

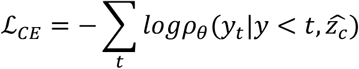

and apply a cell–text alignment loss (InfoNCE) between **z**_*c*_ and the pooled ground-truth caption embedding **z**_*t*_.

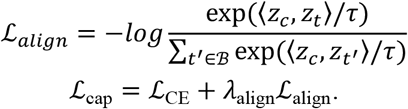

Only the cross-modal projectors, cross-attention, and the language head are trainable. Captions are ontology-grounded to reduce stylistic variance.

### Cell state conditioned pseudo-cell synthesis

To synthesize full macrogene expression vectors that realize a target cellular state described in natural language, a text prompt is encoded by the frozen LLM and pooled to *t*, then projected to the shared latent **z**_*t*_ = *p*_*t*→*z*_(**t**). Metadata *m* modulates *z*_*t*_ to produce 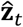. A learned start token represents a generic baseline cell. A regression head with gated cross-attention over 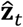 emits a macrogene-level expression vector 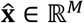 in the fixed orthology-aware vocabulary. Non-negativity and total-count constraints are enforced at output. The reconstruction loss is mean squared error on log-scaled counts:

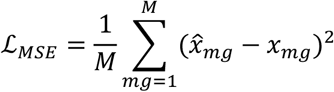

To enforce semantic consistency, a cycle-consistency term re-encodes 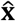 through the frozen encoder and aligns the resulting cell latent 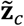 to **z**_*t*_:

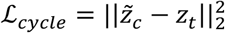

A sparsity regularizer (*l*_1_ on 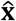) is optionally included to discourage spurious broad activation. The total loss is:

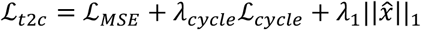

We freeze both the encoder and LLM, and confine trainable parameters to projectors, fusion, and the regression head. We also canonicalized diverse style prompts to standardize conditioning.

### Conditioned perturbation response generation

Given a control cell and a textual specification of diverse perturbations, we aim to generate a post-perturbation expression profile. Specifically, the control expression **x**^ctrl^ is also encoded by the Speciesformer encoder to obtain ***h***_*cell*_ and per-macrogene states 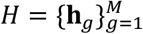. The perturbation description is pooled to **t** and projected to **z**_*t*_, then modulated by **m** to yield 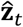. A regression head computes the predicted change Δ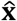 by attending from projected macrogene states {**z**_*g*_} to 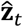 (and, when beneficial, to token-level text keys). The counterfactual is

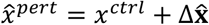

We minimize MSE on log-scaled counts, augmented by (i) a sparsity prior on the delta,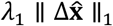, and (ii) an ordinal regularizer when dose or MOI is available, penalizing violations of monotonic ordering across concentrations. When supervision is scarce, we optionally include a consistency term that aligns the encoded latent of 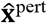 to that of the observed perturbed profile.

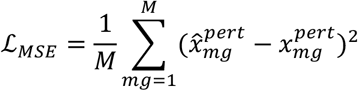

During the main stage, the encoder and LLM are frozen, only projectors, fusion, and the regression head are trained.

### Training setting and optimization

Speciesformer is implemented using PyTorch with distributed data parallel (DDP) training. We employed PyTorch’s automatic mixed precision (AMP) and mixed-precision training with BF16 data type to reduce memory usage and accelerate training.

#### Speciesformer encoder

Pretraining hyperparameters for Speciesformer encoder were optimized by Adam with weight decay fix of 0.01. The learning rate follows linear warm-up for ~5% of total steps to a peak of 1e-4, then cosine decay to a min rate of 6e-5. Unless noted, all embeddings and decoder heads share the model width of 640. Details of hyperparameters for Speciesformer encoder are listed in Supplementary Table 2. The model is pre-trained for 5 epochs under DDP across 16 Ascend 910B1 NPUs. The per-device batch size is 16, with gradient accumulation over 5 minibatches. This configuration sustains stable large-batch optimization while preserving statistical efficiency for masked reconstruction.

#### Speciesformer unified generative decoder

To transform the cross-species encoder from a representation learner into a conditional generative model, we post-trained a unified generative decoder in two consecutive stages. The first stage aimed to establish a shared alignment between cell expression states and textual cell-state descriptions, whereas the second stage specialized the aligned decoder for three generative functions. This two-stage strategy was designed to first anchor transcriptomic and linguistic representations in a common cell-state space, and then optimize task-specific generative objectives without losing the shared biological semantics learned during cross-species pretraining.

During the first post-training stage, we formulated each sample as a paired cell expression-text instance. For each cell, available biological metadata, including species, tissue, cell type, disease state, developmental stage and experimental condition, were converted into a textual description paired with the corresponding expression profile. Transcriptomic and textual representations were aligned using a cross-modal projector containing 32 learnable query tokens, with cross-attention activated every two transformer layers to promote bidirectional information exchange between modalities.

Post-training was performed on 8 NVIDIA A100 GPUs for five epochs. We used the AdamW optimizer with a weight decay of 0.001 and a cosine learning-rate schedule with 5% warm-up. The peak and minimum learning rates were set to 3e-5 and 3e-6, respectively. The batch size was 32 for cell expression and text encoding, and 8 for contrastive learning.

After the unified expression-text alignment stage, the model was further post-trained for three generative functions. Each function was initialized from the stage-1 checkpoint but optimized with its own input format, target output and task-specific loss. This design allowed the three tasks to share a common biological and semantic representation space, while reducing negative transfer among tasks with different output structures. Details for hyperparameters of generative decoder in the stage 2 are listed in Supplementary Table 3.

### Evaluation metrics

We evaluated the model across diverse downstream tasks, including cell type annotation, perturbation prediction, batch integration, cell type description generation, conditional pseudo-cell generation and conditional perturbed-cell generation. For each task, we used metrics that quantify both predictive accuracy and biological fidelity. When applicable, all metrics were computed on held-out test sets, and performance was reported across datasets, species, tissues or perturbation settings. For comparability, we mapped the model output results back to the expression profiles at the gene level before conducting the evaluation in all tasks.

For cell type annotation, we evaluated whether the learned cell embeddings could support accurate cell identity prediction. We used accuracy and macro-averaged F1 score, as the main classification metrics. We employed macro-averaged F1 score because it is used to emphasize performance on rare cell types in the test set.

For cell type description generation, we evaluated whether the model could generate textual descriptions that were both linguistically consistent with the references and biologically faithful to the input expression profiles. We used three complementary groups of metrics. First, we assessed lexical and sequence-level similarity using BLEU(Papineni, et al., 2002), ROUGE(Lin, 2004) and METEOR(Banerjee and Lavie, 2005). BLEU measures n-gram precision, ROUGE captures overlap with reference descriptions, and METEOR provides a more flexible matching criterion that accounts for partial word-level agreement. Second, we evaluated semantic distributional similarity by computing maximum mean discrepancy (MMD) and earth mover’s distance (EMD) between text embeddings of generated and reference descriptions(Xiao, et al., 2024). Third, we assessed biological correctness by extracting cell type labels from the generated descriptions and comparing them with the ground-truth annotations using accuracy and macro-F1 score.

For conditional pseudo-cell generation, we evaluated whether the model could generate realistic gene expression profiles from textual or structured cell-state conditions. At the expression level, we computed Pearson correlation, and Spearman correlation between generated and real expression profiles. To assess biological validity, we evaluated cell type classification accuracy of generated cells using an external or held-out classifier.

For perturbation prediction, we employed Cell-Eval(Adduri, et al., 2025) to evaluate the ability of the model to recover transcriptional responses induced by genetic, chemical or cytokine perturbations. Given a predicted post-perturbation expression profile and the corresponding observed profile, we computed full-expression, perturbation-signature and discrimination level metrics. At the full-expression level, we reported mean squared error (MSE) and mean absolute error (MAE) to quantify global reconstruction error. To evaluate whether the model captured perturbation-induced expression changes, we computed Pearson correlation on expression differences between perturbed and control states, denoted as PCC(Delta), and Spearman correlation on log-fold changes, denoted as LFCSPCC. These metrics assess the direction and relative magnitude of the predicted perturbation responses. To measure recovery of biologically relevant perturbation signals, we evaluated performance on differentially expressed genes. For each perturbation condition, significant DE genes were identified from the observed data using an adjusted P-value threshold, such as adjusted P < 0.05. We then computed correlation on top DE genes, DE-gene overlap (DEOver) and DE precision (DEPrec) to assess whether the predicted profiles recovered the genes most strongly associated with the perturbation. For conditional perturbed-cell generation, we further included perturbation-level discrimination metrics. We adapted the perturbation discrimination score from Wu et al. to evaluate whether generated post-perturbation profiles preserve relative transcriptional differences among perturbation conditions. We report the normalized inverse PDS(L2), where higher values indicate better recovery. We also computed AUROC and AUPRC for DE-gene recovery by assigning binary labels to genes based on significance in the observed data and using predicted significance scores to rank genes. AUROC summarizes the true-positive and false-positive trade-off across thresholds, whereas AUPRC quantifies the precision-recall trade-off and is particularly informative when perturbation-responsive genes are sparse.

