## Supplementary Material for "Speciesformer learns conserved cellular states for cross-species generative virtual cell modeling"

^2^ Independent Researcher

^3^ LC-Bio Technologies (Hangzhou) CO., LTD, Hangzhou, China

**The PDF file includes:**

Supplementary Notes

Supplementary Figure 1 to 12

Supplementary Table 1 to 7

**Supplementary Notes**

1. **Data curation and preprocessing**

We assembled a multi-species scRNA-seq corpus 131M to support pretraining and evaluation of single-cell foundation models. We integrated two sources spanning multiple organisms and assay chemistries:

1.	CELLxGENE Census

We mirrored the 2024-07-01 Census release and its associated per-dataset AnnData/H5AD artifacts via the documented API, covering 115M human and mouse cells and smaller non-mammalian cohorts. We recorded the exact release tag for reproducibility and preserved Census-level integrated covariates without further transformation at the assembly stage.

2.	GEO

To enrich under-represented organisms, we executed automated GEO queries for target taxa, including macaque, marmoset, crab-eating macaque, chimpanzee, gorilla, bonobo, green monkey, rat, chicken. We both included scRNA-seq (dissociated cells) and nuclei assays, downloaded raw count matrices or per-study H5/MTX bundles when available, in total of 56M cells, and captured original multi-level metadata, including samples, species, tissues, genders, development stages, disease states, and library constructions.

Single-cell and single-nuclei RNA sequencing datasets were accepted if labeled as sc/snRNA-seq. We only assembled Public, redistributable datasets with DOIs or accessions. For datasets from CZI CELLxGENE, we used their release manifests. For GEO, we filtered by organism and library strategy (“RNA-Seq”, single-cell). Datasets must expose cell-by-gene count matrices and gene identifiers resolvable to an authoritative gene catalog (Ensembl/NCBI). We checked the schema of datasets from two resources to preliminary validate the data quality. Schema checks included: non-negative integer counts (UMI), consistent feature dimensions, unique cell barcodes, and organism-consistent gene ID namespaces.

We implemented source-agnostic quality control so downstream work can adopt stricter task-specific filters. Specifically, only cells that passed these filtering metrics were used for downstream analyses in this work. Ensembl-annotated protein-coding genes were used for downstream analysis. Cells with less than 200 detected Ensembl-annotated protein-coding genes were excluded. We performed library-size normalization and log1p transformation on raw counts for each dataset.

2. **Macrogene construction**

When modelling single-cell datasets from multiple species, a central challenge is how to identify homologous relationships among genes across lineages. A common strategy is to integrate cross-species transcriptomes by mapping one-to-one orthologous genes between two species. However, this pairwise ortholog-based approach does not scale well to the multi-species setting considered in our study. Instead, we employed the kernel archetypal analysis(Cutler and Breiman, 1994) to construct a shared “macrogene” space at the cross-species level. Briefly, we coupled multi-species single-cell transcriptomes with protein language model representations to learn a set of cross-species macrogenes in a protein embedding space. For each species, we first collected the protein sequences corresponding to all genes and computed protein embeddings using a pre-trained ESM2 protein language model(Lin, et al., 2023). For genes with multiple protein products, we averaged the respective embedding vectors to obtain a unified protein embedding matrix P covering all genes across species. In this embedding space, we then constructed a k-nearest-neighbor graph based on Euclidean distances between protein embeddings and, using a manifold-based model, derives a non-negative “gene-macrogene” weight matrix $W$. This ensures that genes that are close in protein functional space are grouped into highly similar gene clusters, thereby yielding a stable macrogene space.

To capture gene homology across different taxonomic levels, we further constructed macrogenes by progressively extracting gene homology according to the evolutionary relationships encoded in the phylogenetic tree. For genes drawn from different classes, phyla, families and species, we first extracts macrogenes within groups sharing the same taxonomic attribute and then iteratively aggregates them. For example, for human, chimpanzee, bonobo and gorilla, we first derived Hominidae family-level macrogenes from genes in these four species. In a second iteration, we then extracted macrogenes from three sets of such family-level macrogenes. Using this inverse phylogenetic tree strategy, we obtained macrogenes layer by layer from species with shared attributes. Ultimately, we extracted 862 macrogenes from 11 species. Using the learned gene-to-macrogene mapping weights, we converted the single-cell expression matrices of all 11 species into a unified cell-by-macrogene representation for pretraining and downstream analyses.

Macrogene sequencing embeddings

For each gene, we retrieved the canonical DNA and protein sequences from Ensembl(Dyer, et al., 2024) and the curated textual description from NCBI Gene. When an NCBI description was unavailable, a short scientific synopsis was generated using ChatGPT API with a constrained prompt to avoid unverifiable claims. The generated text was stored alongside the provenance flag (“LLM-derived”) for downstream auditing.

Three pretrained encoders were used to derive modality-specific embeddings per gene: (i) a Nucleotide Transformer for genomic DNA, (ii) ESM-2b for amino-acid sequences, and (iii) Llama-3 for short free-text descriptions. For sequence inputs, the model-recommended tokenization was applied. Sequences exceeding the model context were truncated at the 5′ end after prioritizing the principal isoform, and sequences shorter than the context window were left unpadded. Embedding vectors were obtained by mean-pooling the final hidden states over tokens (or using the model’s [CLS] representation, when recommended in its documentation) and L2-normalized. Text inputs were lower-cased and stripped of citations, accession numbers, and markup prior to encoding.

**Supplementary Figures**

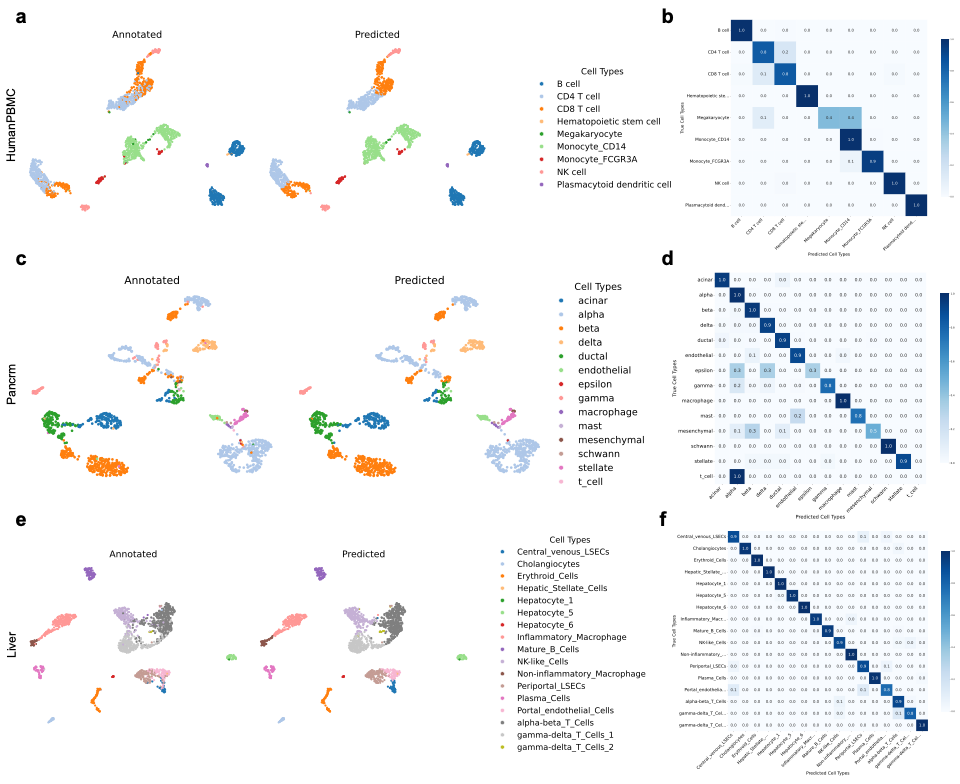

**Supplementary Figure 1**. Cell type annotation results. The UMAP of cell embeddings and confusion matrices from Speciesformer on HumanPBMC (**a, b**), Pancrm (**c, d**), and Liver (**e, f**) datasets, respectively.

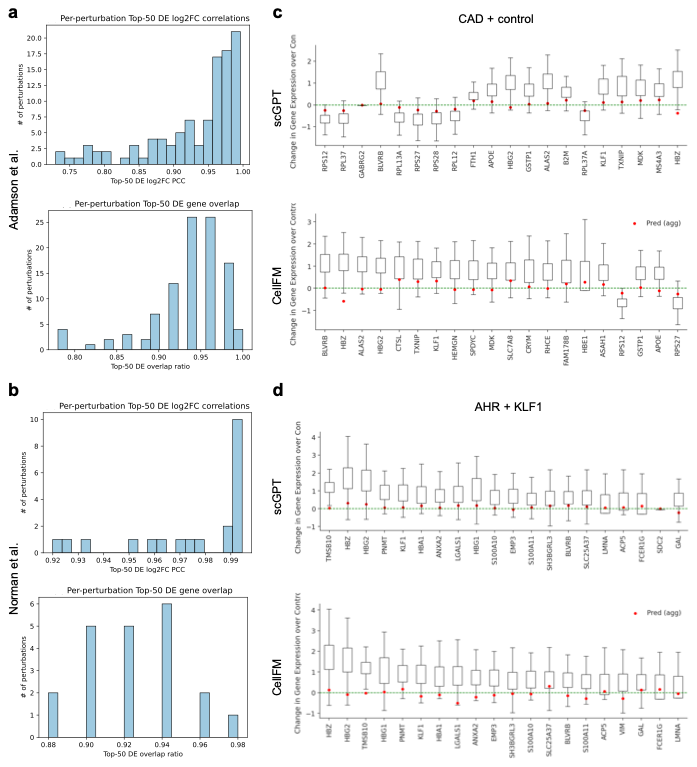

**Supplementary Figure 2**. Perturbation prediction analysis. Distribution of per-perturbation prediction at Top-50 differentially expressed genes from Speciesformer on (**a**) Adamson et al. and (**b**) Norman et al. datasets. Example perturbations of scGPT and CellFM in the (**c**) Adamson test dataset (single gene perturbation: CAD), and Norman test dataset (perturbed combination: AHR + KLF1), distribution of predicted and actual gene expression change of the top 20 differentially expressed genes. The box denotes the interquartile range of expression change. The median is marked by the central line within each box. Whiskers extend to 1.5 times the interquartile range. Horizontal dashed lines represent the null baseline of gene expression changes.

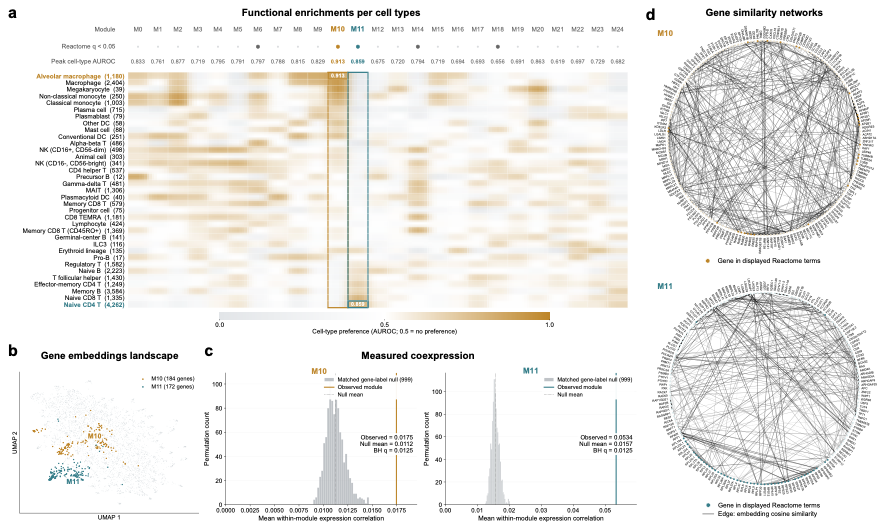

**Supplementary Figure 3**. Cell type-specific activated gene programs extracted from Speciesformer’s embeddings in the immune human dataset. Coloring indicates average gene expression. For gene programs containing more than 10 genes (indicated by an asterisk), the first 10 genes are shown for simplicity.

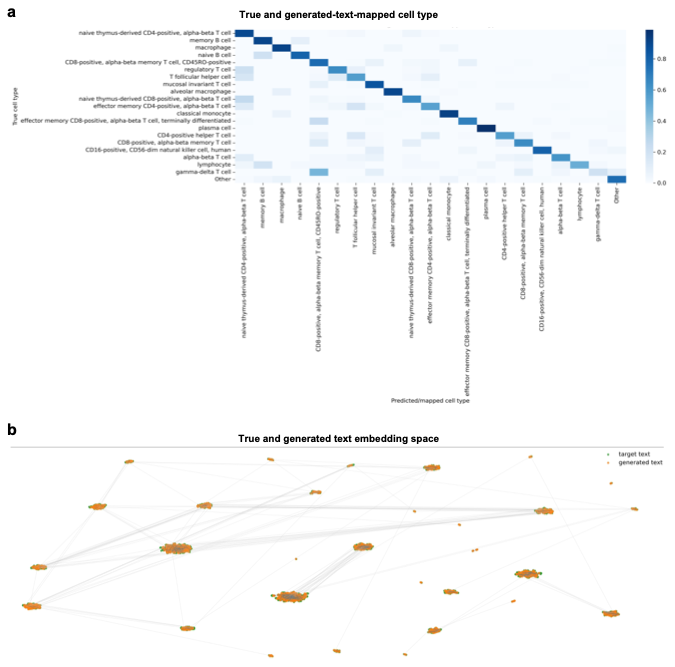

**Supplementary Figure 4**. **a**, Confusion matrices of Immune dataset between true cell type labels and generated cell type descriptions mapped cell type labels from Speciesformer. **b**, UMAP of target text embeddings and generated text embeddings.

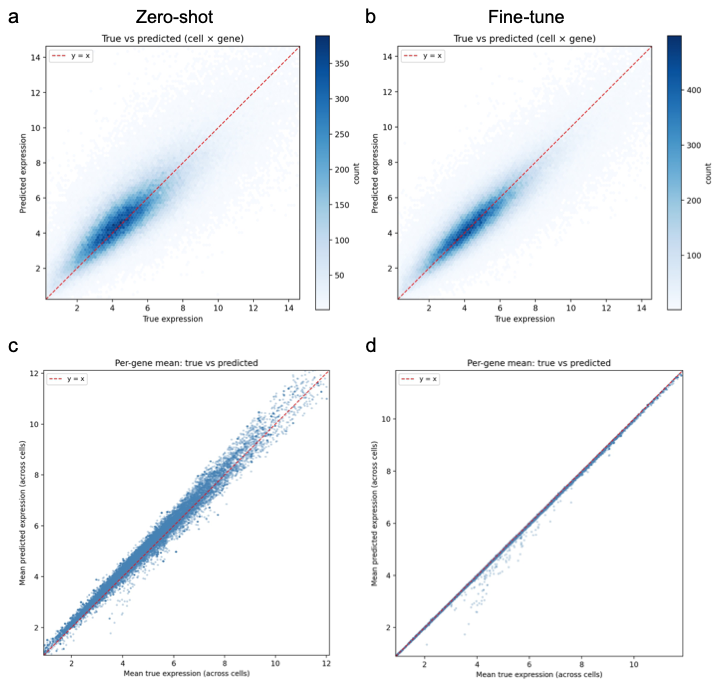

**Supplementary Figure 5**. Hexbin and per-gene scatter plots on Immune dataset between true and predicted gene expression under zero-shot (**a, c**) and fine-tune setting (**b, d**).

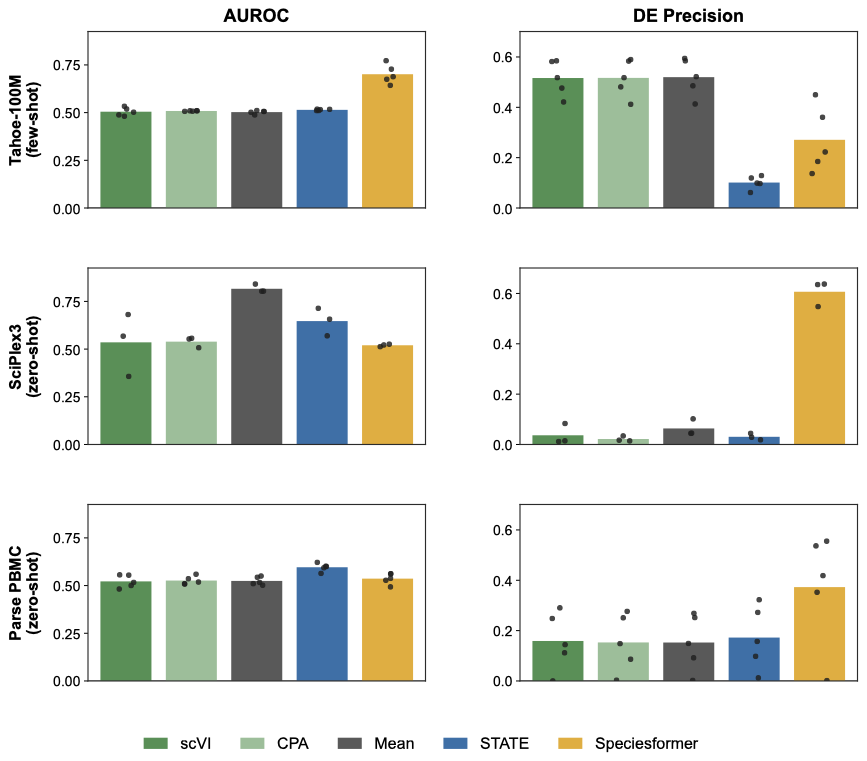

**Supplementary Figure 6**. Additional metrics for perturbation-conditioned cellular response prediction performance across three perturbation datasets (Chemical perturbation, Tahoe-100M; Multiplex compounds screen, sciplex3; Cytokines perturbation, Parse-PBMC). Comparisons included the mean baseline, two autoencoder-based models scVI and CPA, and a transformer-based model design for perturbation response prediction STATE. (Left) Area Under the Receiver Operating Characteristic curve (AUROC), summarizing model performance in identifying true DE genes. (Right) DE genes precision, measuring the predicted DE genes that were truly significant in the observed data.

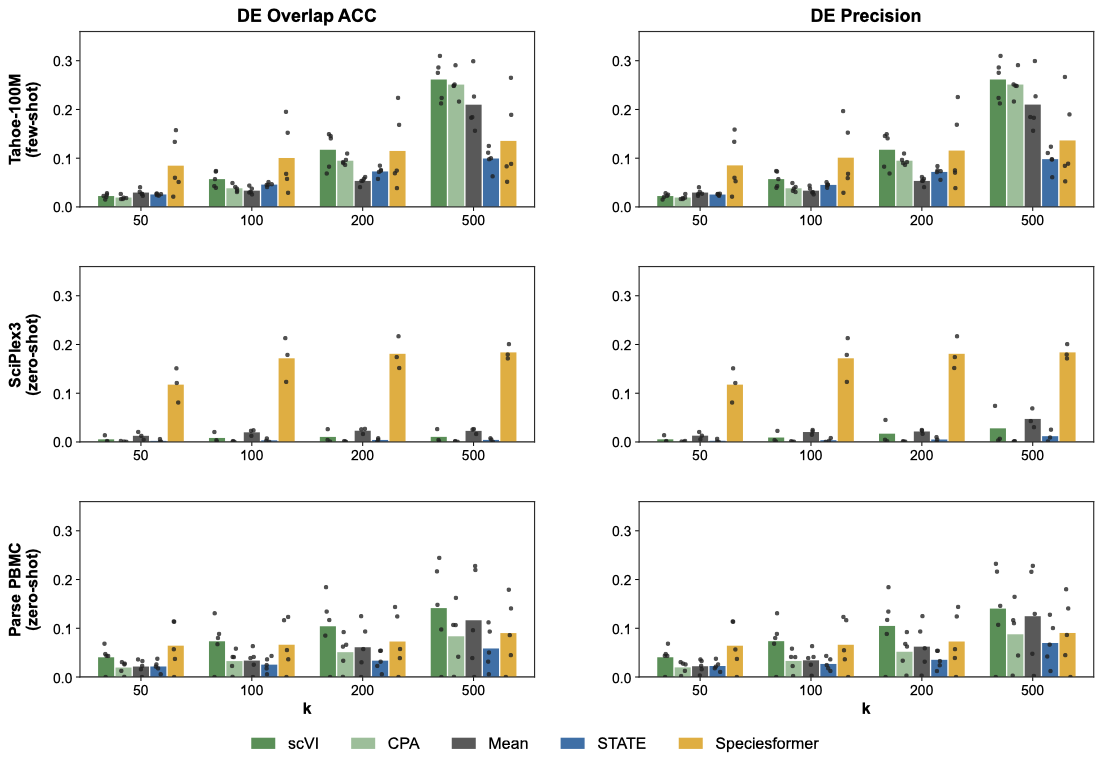

**Supplementary Figure 7**. Perturbation response prediction for top differentially expressed (DE) genes. (Left) DE genes Overlap accuracy, accessing fraction of top DE genes in the observed data that were also identified in model predictions. (Right) DE genes precision, measuring the predicted DE genes that were truly significant in the observed data. Two metrics were evaluated across various values of k (50, 100, 200, 500).

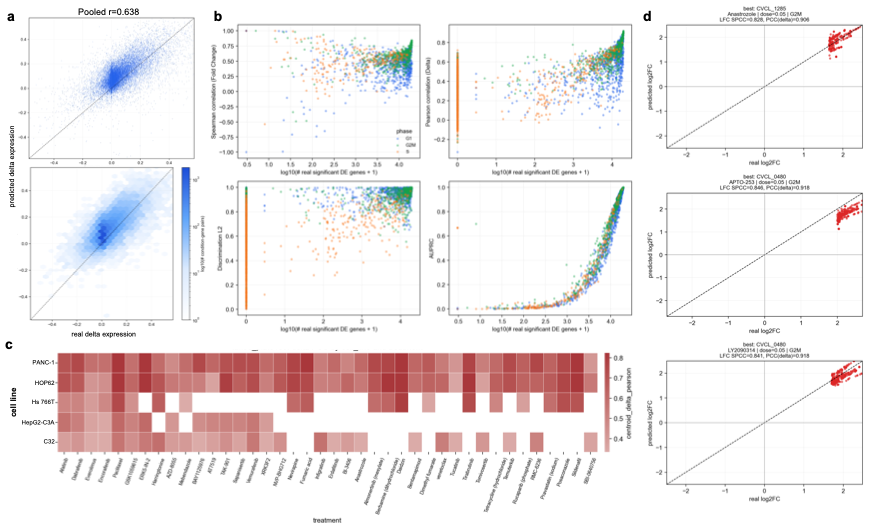

**Supplementary Figure 8**. Post-perturbation delta expression prediction by Speciesformer across cell-cycle phase, cell lines, treatments, and doses of Tahoe-100M. a, Scatter (up) and hexbin (down) plots of overall delta gene expression prediction. b, Performance across perturbation-response strengths. Each point represents a perturbation condition, with response strength quantified by the number of true significant DE genes. The four panels report Spearman correlation of log-fold changes, Pearson correlation of gene-expression changes (Delta), discrimination L2, and AUPRC, stratified by cell-cycle phase. c, Cross-context recovery of drug-induced expression changes. The heatmap summarizes gene-wise Pearson correlations between true and predicted expression changes (Delta) for each cell line–treatment pair. Blank cells represent cell line-treatment combinations without an evaluable condition. d, Recovery of drug-induced transcriptional effects in three best-performing perturbation contexts. Each panel compares true and predicted log2 fold changes for the top response-associated genes in one perturbation condition.

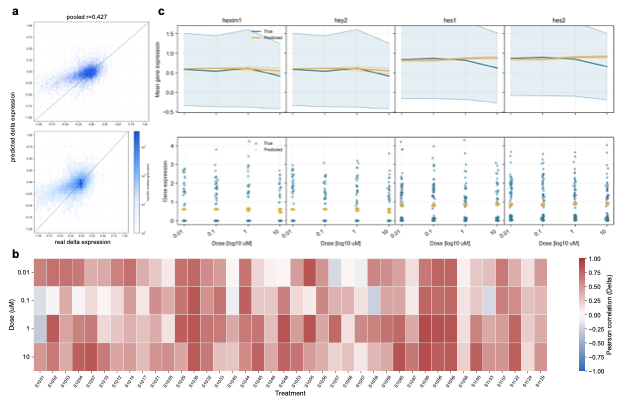

**Supplementary Figure 9**. Speciesformer captures condition-specific and dose-dependent perturbation responses in sciPlex3 MCF7 cells. a, Scatter (up) and hexbin (down) views of true and predicted gene-expression changes (Delta) across pooled treatment-dose conditions. For hexbin panel, color from white to blue indicates increasing log10 density of condition-gene pairs. b, Dose-stratified perturbation-response recovery. The heatmap shows gene-wise Pearson correlations between true and predicted expression changes (Delta) across four doses for 40 drug treatments. c, Dose-dependent expression responses to the unseen drug S2818. True and predicted expression profiles are shown across four doses for representative differentially expressed genes, illustrating recovery of both dose-dependent mean responses and single-cell expression distributions.

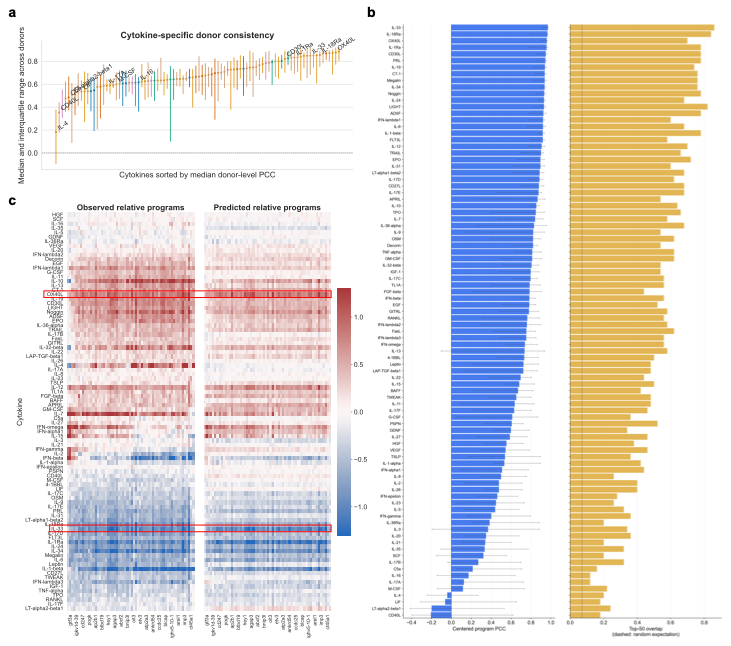

**Supplementary Figure 10**. Speciesformer accurately identified cytokine response programs in held-out CD4 memory T cells. a, Donor-level consistency of cytokine response-program recovery. Points show median observed-predicted centred program PCC across 12 donors, with interquartile ranges. Cytokines are ordered by median centred program PCC. b, Performance of predicted donor-centred cytokine response programs. Cytokines are ranked by centred program PCC (left; 95% donor-bootstrap confidence intervals); Top-50 feature overlap is shown at right. Dashed line, random overlap expectation (50/709). c, Model recovery of cytokine-specific relative response programs. Donor-centred programs are shown for observed data and model predictions. Cytokines and features are ordered using the observed programs.

**Supplementary Tables**

**Supplementary Table 1**. Pre-trainning data compositions and sources of Speciesformer across 11 species.

| Species | Number of cells | Source |
| --- | --- | --- |
| Homo sapiens | 74M | CxG |
| Mus musculus | 41M | CxG |
| Macaca mulatta | 7M | CxG & GEO |
| Rattus norvegicus | 3M | GEO |
| Callithrix jacchus | 2M | CxG & GEO |
| Macaca fascicularis | 1.5M | GEO |
| Pan troglodytes | 1.5M | CxG & GEO |
| Gallus gallus | 1M | GEO |
| Chlorocebus sabaeus | 0.2M | GEO |
| Gorrila gorrila | 0.2M | CxG & GEO |
| Pan panisus | 0.1M | GEO |

**Supplementary Table 2**. Hyperparameters for two Speciesformer encoders with different size.

| Hyperparameter | 100M | 500M |
| --- | --- | --- |
| input length | 862 | 862 |
| masking ratio | 15% | 15% |
| # of encoder blocks | 12 | 24 |
| # of attention heads | 10 | 20 |
| hidden size | 640 | 1280 |
| hidden size for cell tokens | 128 | 128 |
| learning rate | 1e-4 | 1e-5 |
| scheduler | cosine annealing | cosine annealing |
| weight decay | 1e-2 | 1e-1 |
| batch size | 32 | 32 |
| # of gradient accumulation | 8 | 8 |
| # of GPU | 16 | 16 |
| GPU type | Ascend 910 NPU | Ascend 910 NPU |
| # of epochs | 5 | 10 |
| warmup ratio | 5% | 5% |

**Supplementary** **Table 3**. Hyperparameters for the unified generative decoder of Speciesformer

| Hyperparameter | Cell-type description generation | Conditional pseudo-cell generation | Conditional perturbed-cell generation |
| --- | --- | --- | --- |
| max length for text | 128 | 256 | 256 |
| learning rate | 3e-5 | 5e-5 | 1e-5 |
| mini learning rate | 3e-6 | 5e-6 | 1e-6 |
| scheduler | cosine annealing | cosine annealing | cosine annealing |
| weight decay | 1e-3 | 1e-3 | 1e-3 |
| batch size | 16 | 32 | 16 |
| # of gradient accumulation | 16 | 8 | 16 |
| # of GPU | 8 | 8 | 8 |
| GPU type | A100 | A100 | A100 |
| # of epochs | 2 | 2 | 2 |
| warmup ratio | 5% | 5% | 5% |

**Supplementary Table 4**. Benchmarking results from Speciesformer on cell type annotation task. We benchmarked Speciesformer against to four single-cell foundation models Geneformer, scGPT, UCE, CellFM across five datasets, including three intra-datasets HumanPBMC, Immune, and Pancrm, and two inter-datasets human pancreas (hPancreas) and Liver. Two classification evaluation metrics are present in this benchmarking results.

| Models | HumanPBMC | Immune | Pancrm | hPancreas | Liver |
| --- | --- | --- | --- | --- | --- |
| Accuracy | | | | | |
| scGPT | 0.618 | 0.621 | 0.556 | 0.763 | 0.521 |
| Geneformer | 0.608 | 0.473 | 0.828 | 0.949 | 0.62 |
| UCE | 0.96 | 0.918 | 0.841 | 0.885 | **0.897** |
| CellFM | **0.966** | 0.925 | 0.962 | 0.978 | 0.67 |
| Speciesformer | 0.91 | **0.93** | **0.971** | **0.984** | 0.776 |
| Macro-F1 | | | | | |
| scGPT | 0.244 | 0.27 | 0.209 | 0.559 | 0.236 |
| Geneformer | 0.523 | 0.675 | 0.455 | **0.864** | 0.305 |
| UCE | 0.886 | 0.884 | 0.53 | 0.749 | 0.604 |
| CellFM | 0.775 | 0.888 | 0.727 | 0.699 | 0.632 |
| Speciesformer | **0.901** | **0.899** | **0.873** | 0.751 | **0.659** |

**Supplementary Table 5**. Benchmarking results from Speciesformer on perturbation response prediction task. We benchmarked Speciesformer against to two single-cell foundation models scGPT, and CellFM across two genetic perturb-seq datasets: Adamson et al. and Norman et al. Six classification evaluation metrics are present in this benchmarking results.

| Models | PCC (Delta) | PCC DE | LFC-SPCC | AUPRC | AUROC | DEO | DEP | MSE DE | PDS |
| --- | --- | --- | --- | --- | --- | --- | --- | --- | --- |
| Adamson et al. | | | | | | | | |  |
| scGPT | 0.618 | 0.663 | 0.533 | 0.544 | 0.469 | 0.409 | 0.43 | 0.26 | 0.524 |
| CellFM | **0.71** | 0.717 | 0.55 | 0.488 | 0.453 | 0.303 | 0.394 | 0.32 | 0.519 |
| Speciesformer | 0.98 | **0.99** | **0.99** | **0.981** | **0.766** | **0.999** | **0.998** | **0.1** | **0.524** |
| Norman et al. | | | | | | | | | |
| scGPT | 0.49 | 0.49 | 0.601 | 0.209 | 0.413 | 0.025 | 0.121 | 0.26 | 0.505 |
| CellFM | 0.385 | 0.385 | 0.271 | 0.207 | 0.398 | 0.032 | 0.13 | 0.32 | **0.506** |
| Speciesformer | **0.99** | **0.989** | **0.998** | **0.985** | **0.827** | **0.996** | **0.996** | **0.08** | 0.505 |

**Supplementary Table 6**. Benchmarking results of cell description generation on immune dataset. Speciesformer was benchmarked with three multimodal foundation models, GPT2, C2S (Large), and scMMGPT. We present seven text generation evaluation metrics: Accuracy, Macro-F1, BLEU-2, ROUGE-2, METEOR, maximum mean discrepancy (MMD) and earth mover’s distance (EMD).

| Model | Acc | Macro-F1 | BLEU-2 | ROUGE-2 | METEOR | MMD | EMD |
| --- | --- | --- | --- | --- | --- | --- | --- |
| GPT2 | 0.339 | 0.16 | 0.413 | 0.352 | 0.44 | 0.127 | 0.02 |
| C2S (large) | 0.592 | 0.55 | 0.734 | 0.685 | 0.743 | 0.01 | 0.005 |
| scMMGPT | 0.692 | 0.38 | 0.66 | 0.65 | 0.685 | 0.024 | 0.0045 |
| Speciesformer | **0.773** | **0.724** | **0.856** | **0.823** | **0.86** | **0.001** | **0.0004** |

**Supplementary Table 7**. Cell-type transfer evaluation between real cells and generated pseudo-cells. Speciesformer is compared with generative or multimodal models, including scVI, scDiffusion, scGPT, C2S, scMMGPT and InstructCell.

| Models | Top 3 | Top 5 | Top 10 | Top 25 |
| --- | --- | --- | --- | --- |
| scGPT | 0.184 | 0.179 | 0.181 | 0.188 |
| scVI | 0.244 | 0.24 | 0.243 | 0.235 |
| scDiffusion | 0.234 | 0.229 | 0.237 | 0.235 |
| C2S | 0.259 | 0.257 | 0.275 | 0.272 |
| scMMGPT | 0.3 | 0.299 | 0.299 | 0.298 |
| InstructCell | 0.376 | 0.392 | 0.39 | 0.31 |
| Speciesformer-Zeroshot | 0.643 | 0.637 | 0.733 | 0.668 |
| Speciesformer | **0.651** | **0.653** | **0.764** | **0.71** |
